# The structure of pathogenic huntingtin exon-1 defines the bases of its aggregation propensity

**DOI:** 10.1101/2022.10.25.513661

**Authors:** Carlos A. Elena-Real, Amin Sagar, Annika Urbanek, Matija Popovic, Anna Morató, Alejandro Estaña, Aurélie Fournet, Xamuel L. Lund, Zhen-Dan Shi, Luca Costa, Aurélien Thureau, Frédéric Allemand, Rolf E. Swenson, Pierre-Emmanuel Milhiet, Alessandro Barducci, Juan Cortés, Davy Sinnaeve, Nathalie Sibille, Pau Bernadó

## Abstract

Huntington’s Disease is a neurodegenerative disorder caused by a CAG expansion of the first exon of the *HTT* gene, resulting in an extended poly-glutamine (poly-Q) tract in the N-terminus of the protein huntingtin (httex1). The structural changes occurring to the poly-Q when increasing its length remain poorly understood mainly due to its intrinsic flexibility and the strong compositional bias of the protein. The systematic application of site-specific isotopic labeling has enabled residue-specific NMR investigations of the poly-Q tract of pathogenic httex1 variants with 46 and 66 consecutive glutamines. The integrative analysis of the data reveals that the poly-Q tract adopts long α-helical conformations stabilized by glutamine side-chain to backbone hydrogen bonds. ^19^F-NMR of site-specifically incorporated fluoro-glutamines and molecular dynamics simulations demonstrate that the mechanism propagating α-helical conformations towards the poly-Q from the upstream N17 domain is independent of the poly-Q track length. Aggregation and atomic force microscopy experiments show that the presence of long and persistent α-helices in the poly-Q tract is a stronger signature in defining the aggregation kinetics and the structure of the resulting fibrils than the number of glutamines. The ensemble of our observations provides a structural perspective of the pathogenicity of expanded httex1 and paves the way to a deeper understanding of poly-Q related diseases.

## Introduction

Among the nine neurodegenerative disorders caused by expansions of polyglutamine (poly-Q) tracts, Huntington’s Disease (HD) stands out due to its prevalence and devastating effects^1^. HD is triggered by an abnormal expansion of the poly-Q tract located in exon1 (httex1) of the 348-kDa huntingtin, a ubiquitous protein involved in multiple pathways^2,3^. In its non-pathogenic form, the httex1 poly-Q tract is comprised of 17-20 glutamines^4^; however, when the number of consecutive glutamines exceeds the pathogenic threshold of 35, it results in aggregation-prone mutants. Indeed, fragments of mutant httex1 can be found forming large cytoplasmic and nuclear aggregates within neurons of the striatum, a well-known hallmark of HD^5,6^. The presence of such aggregates, the neuronal degeneration, the age of onset and disease severity, all correlate with the length of the expanded poly-Q tract^7^. Notably, the mutant httex1 fragment alone suffices to reproduce HD symptoms in mice^8^. Unfortunately, no effective treatment is currently available, mainly because of the lack of knowledge regarding the molecular mechanisms underlying the disease^9^.

Amyloidogenic aggregates were initially proposed to be the toxic species in HD, acting by sequestering essential cellular proteins and components^10^. However, they have also been related to neuronal survival in certain cell types, suggesting that aggregates may be protective^11–14^. It has also been proposed that toxicity is caused by soluble species of httex1, which are characterized by a high polymorphism comprising monomers, dimers, tetramers and small oligomers^15,16^. It has been shown that N17, the 17 residue-long fragment preceding poly-Q (Fig. 1a), is the pivotal element triggering httex1 self-recognition and enhancing aggregation^17–19^. However, the structural mechanism by which N17 propagates aggregation and cytotoxicity of expanded httex1 remains to be defined.

**Figure 1.**
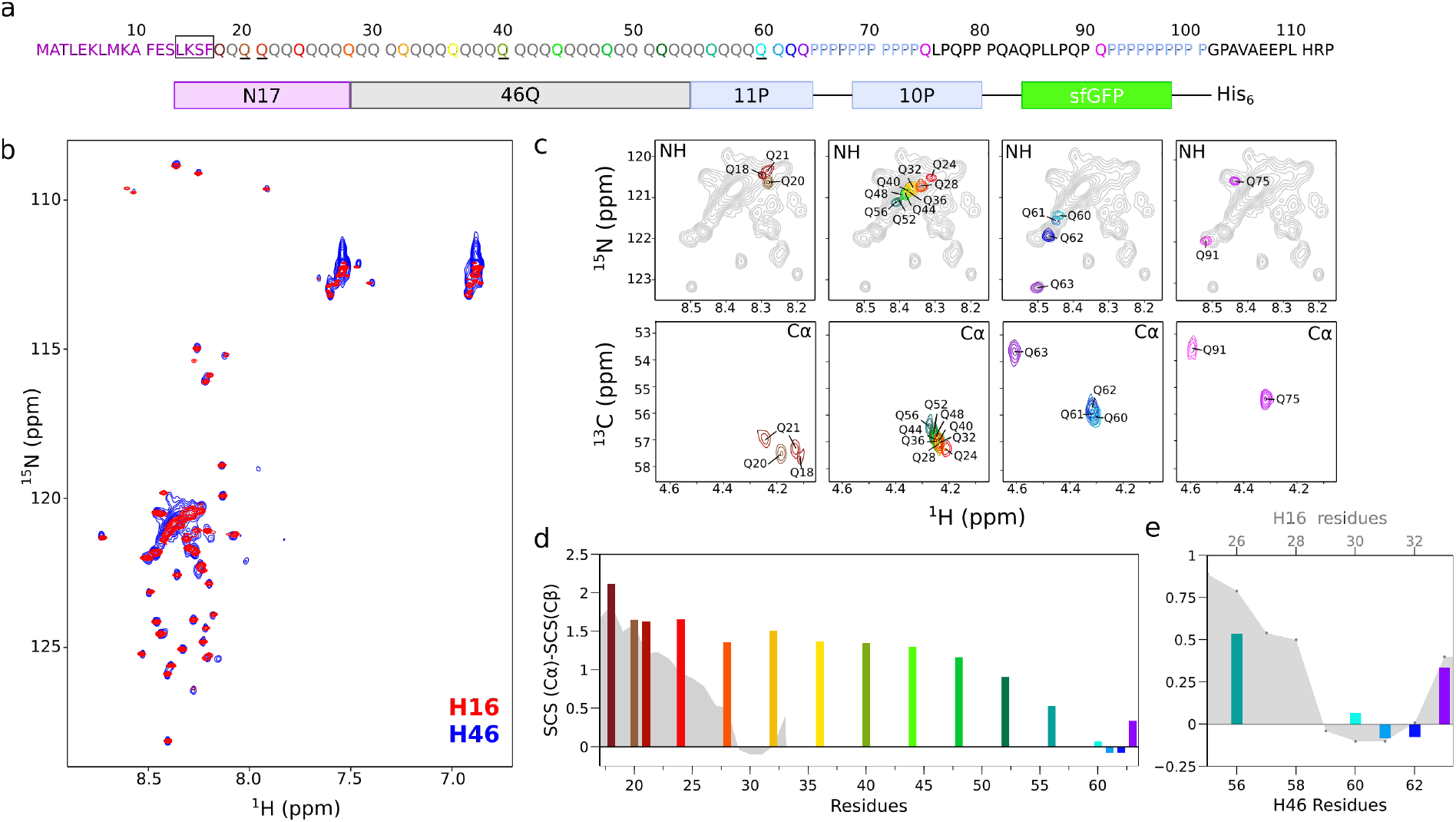
NMR analyses of H46 and comparison with H16. (**a**) Primary sequence of H46 and scheme of the sfGFP-fused construct used in this study. The color code identifies the site-specifically labeled glutamines throughout the study. Underlined glutamines indicate the positions in which 2S,4R-fluoroglutamine (4F-Gln) was introduced for ^19^F-NMR experiments. The box indicates the section mutated in the LKGG-H46 and LLLF-H46 mutants. (**b**) Overlay of the ^15^N-HSQC spectra of fully labeled H46 (blue) with the previously reported for H16 (red)^33^. (**c**) Zoom of the ^15^N-HSQC (upper panels) and ^13^C-HSQC (lower panels) with individually colored SSIL spectra showing the poly-Q NH and Cα regions for different glutamine clusters (Q18-Q21; Q24-Q56; Q60-Q63; and PRR glutamines). In the upper panels, the ^15^N-HSQC of the fully labeled ^15^N-H46 sample is shown in gray. (**d**) Secondary chemical shift (SCS) analysis of H46 poly-Q using experimental Cα and Cβ chemical shifts and a neighbor-corrected random-coil library^37^. The SCS analysis of H16 is shown in gray^33^. (**e**) Comparison of the SCS values of glutamines flanking the PRR in H16 (gray area) and H46 (colored bars) when aligning both sequences from the C-terminus of the poly-Q tract.

Two models have been suggested to connect the pathological threshold and toxicity, the ‘toxic structure’ and the ‘linear lattice’ models^20^. While the ‘toxic structure’ model proposes the appearance of a distinct toxic conformation when the poly-Q tract is expanded beyond the pathological threshold^21,22^; the ‘linear lattice’ model suggests that poly-Q tracts are inherently toxic and that their toxicity systematically increases with the homorepeat length^23,24^. Intriguingly, both models have been supported by antibody recognition experiments in different studies. Furthermore, the absence of sharp changes in single-molecule Förster resonance energy transfer (smFRET), circular dichroism and electron paramagnetic resonance (EPR) experiments around the pathological threshold has been argued to substantiate the ‘linear lattice’ model^25–27^. High-resolution structures of non-pathogenic and pathogenic httex1 variants are required to evaluate the changes occurring upon poly-Q expansion, discriminate between both models and finally define the bases of httex1cytotoxicity.

Until recently, the detailed high-resolution structural characterization of soluble httex1, especially of those with pathogenic poly-Q length, has been hampered by the intrinsic properties of the protein, namely the highly flexible nature, its aggregation propensity and the strong compositional bias. X-ray diffraction or electron microscopy, normally used to study folded proteins, cannot be applied to probe disordered proteins such as httex1. Furthermore, the presence of low complexity regions, such as the long glutamine and proline homorepeats, results in important signal overlap when nuclear magnetic resonance (NMR) is used, hampering the collection of high-resolution information^28^. Despite these difficulties, several high-resolution NMR studies of non-pathogenic httex1 and N-terminal fragments have been reported^19,26,27,29,30^. However, due to the severe overlap, only assignments of the first and last glutamines of the poly-Q tract could be achieved.

To circumvent the NMR signal overlap, our lab recently developed a site-specific isotopic labeling (SSIL) strategy that combines cell-free protein expression and non-sense suppression, enabling the investigation of homorepeats in a residue-specific manner^31,32^. When applying this methodology to a non-pathogenic version of httex1 containing a 16-residue-long poly-Q tract (H16), it was shown that the protein was enriched in helical conformations whose length and stability were defined by flanking regions^33^. While the helical propensity was propagated from N17 to the poly-Q through a hydrogen-bond network, it was blocked by the helix-breaking effect caused by the proline-rich region (PRR) that follows the poly-Q tract (Fig. 1a). The structural effects imposed by N17 and PRR may explain the positive and negative regulation of httex1 aggregation by both poly-Q flanking regions, respectively^34,35^. Whether these structural mechanisms also govern the conformational landscape of pathogenic httex1 remains to be discerned.

In the present study, we applied SSIL to a pathogenic form of httex1 containing a 46 residue-long poly-Q tract (H46) to unambiguously assign sixteen of these glutamines spread along the tract and probed the structure and dynamics of the homorepeat. The integration of the NMR information and small-angle X-ray scattering (SAXS) data provided the structural description of H46 as an ensemble of elongated, partially helical conformations, whose propagation and stability mechanisms were deciphered by ^19^F-NMR and molecular dynamics (MD) simulations. All together, our observations provides a detailed structural perspective of the ‘linear lattice’ toxicity model, demonstrating that the presence of long, persistent, aggregation-prone α-helices is concomitant to the expansion of the poly-Q tract beyond the pathological threshold.

## Results

### Pathogenic and non-pathogenic httex1 forms present similar structural features

In order to study a pathogenic form of httex1, a construct comprising the N17 domain, a 46-residue-long poly-Q tract and the PRR was fused to superfolder GFP (sfGFP) (Fig. 1a). A fully ^15^N-labeled H46 sample was prepared and a ^15^N-HSQC NMR spectrum was recorded in order to evaluate its general spectroscopic features. Similarly to H16^33^, the spectrum revealed that while peaks from N17 and the PRR were well dispersed, the peaks corresponding to glutamine residues remained in a large poorly resolved density (Fig. 1b). As expected, the increased molecular weight led to a general peak broadening, which was particularly striking in the region of the expanded poly-Q tract.

The reduced stability of H46 at high concentration precluded the use of traditional 3D-NMR experiments for the frequency assignment. Thus, to confirm the similarities found between the pathogenic and non-pathogenic httex1 constructs, selectively labeled samples of H46 were prepared (^15^N-Ala and ^15^N-Lys; ^15^N-Gly, ^15^N-Ser and ^15^N-Arg; ^15^N-Leu and ^15^N-Glu; and ^15^N-Phe), therefore reducing the overlap of some N17 and PRR signals (Fig. S1a). The selective labeling showed that the vast majority of peaks corresponding to N17 and PRR residues nicely overlap for both H46 and H16. Interestingly, F17 peaks in H16 and H46 displayed different chemical shifts. Altogether, our observations suggest that both httex1 forms share similar structural features outside of the poly-Q tract, although some perturbations in the N17/poly-Q boundary are induced upon the extension of the homorepeat.

In order to obtain high-resolution information on the poly-Q tract of H46, the SSIL approach using previously optimized protocols^36^ was used to specifically study 16 of the 46 glutamines of the H46 poly-Q tract, as well as two glutamines within the PRR (Fig. 1a). ^15^N-HSQC spectra of these samples revealed that the glutamines adjacent to N17 (Q18, Q20 and Q21) adopted the lowest chemical shift values of the poly-Q region without any specific trend, while the following glutamines (Q24-Q56) exhibited steadily increasing ^1^H and ^15^N chemical shifts. This last observation points towards a gradual structural change along the homorepeat (Fig. 1c, upper panels). Finally, the last glutamines of the tract (Q61-Q63) were found at the highest chemical shifts, with Q63, affected by the adjacent poly-P tract, displaying an isolated peak outside of the poly-Q density. The signals corresponding to Q75 and Q91 in the PRR did not show any specific trend. The same features were observed when monitoring the Cα-Hα correlations of the same SSIL samples (Fig. 1c, lower panels). Notably, the Q21 SSIL sample presented two Cα peaks with similar intensity, suggesting the presence of two slowly interconverting conformations. While one of the peaks appeared close to these of Q18 and Q20, the second one was shifted to the same spectral region of the central glutamines of the homorepeat.

The secondary chemical shift (SCS) analysis of the glutamines using a neighbor-corrected random coil database^37^ showed that the poly-Q tract was highly enriched in α-helical conformations, in line with previous observations for non-pathological constructs^26,29,33^ (Fig. 1d). However, this propensity was not homogeneous; the helicity reached its maximum at Q18 and presented a steady plateau over 30 residues, until around Q48, in which the helical propensity was maintained. Only for the C-terminal part of the H46 poly-Q a smooth decrease of helicity was observed, reaching negative SCS values for the last glutamines of the tract. When comparing this SCS analysis with that previously reported for H16^33^, an increase in helicity for the N-terminal part of the H46 poly-Q tract was observed. This phenomenon also explains the shift of the NH, NεH_2_ and CαHα NMR signals of Q18 and Q21 towards more helical positions in H46 when compared with H16 (Fig. S1b). Interestingly, the two Cα peak observed for Q21 presented SCS values of 1.27 and 1.63 ppm, corresponding to two conformations with different helical content.

The extent of the structural effects induced by the PRR was analyzed by aligning the SCS values of H16 and H46 from the C-terminus of the poly-Q tract (Fig. 1e). Interestingly, while the last four glutamines of the tract (Q30-Q33 in H16 and Q60-Q63 in H46) showed the same conformational trends, this similarity was reduced for more distant glutamines, such as Q56. Notice that this residue is closer to the helix-promoting N17 in H16 (Q26) than in H46.

### H66 substantiates the persistence of long α-helical conformations in long poly-Q tracts

The above results suggest that the distance to the PRR is the only parameter defining the length of α-helical conformations in httex1 and that helical conformations encompass larger sections of the homorepeat when the length of the poly-Q tract is increased. In order to validate this hypothesis, we studied an httex1 construct with 66 consecutive glutamines (H66). The ^15^N-HSQC of H66 produced by cell-free expression nicely overlapped with that of H46, indicating that no structural changes occur in httex1 upon incorporating twenty additional glutamines (Fig. S2a). The glutamine region of H66 and H46 spectra displayed the same elongated shape and their maximum intensities were centered at the same proton and nitrogen frequencies. Then, using our standard protocols, we applied the SSIL strategy to two glutamines (Q56 and Q76) of H66 to probe their helical content. Note that these two residues are located at equivalent positions to residue Q56 in H46 when aligning both sequences from the N- and C-termini, respectively (Fig. 2a). The NH peak of H66-Q76 appeared in a position equivalent to H46-Q56, whereas H66-Q56 had a lower chemical shift, suggesting that this glutamine adopted a more helical structure than the other two residues (Fig. 2b). This observation was further confirmed by monitoring the Cα-Hα correlations for the three residues (Fig. 2c). Indeed, while the Cα-Hα peaks for H46-Q56 and H66-Q76 overlapped, H66-Q56 was shifted towards more helical conformations. The helical propensity of these residues was quantified with the SCS analysis, indicating an enhanced helicity for H66-Q56 (1.18 ppm) when compared with H66-Q76 (0.50 ppm) and H46-Q56 (0.53 ppm) (Fig. 3c). Interestingly, the SCS value for H66-Q56 was very similar to those observed for the plateau in H46, indicating that the additional twenty glutamines in H66 adopt helical conformations (Fig. S2b). These results evidenced that the extent of the α-helix-breaking capacity of the PRR is the same in all httex1 forms, allowing the α-helical propensity to be propagated through a larger number of glutamines when the length of the homorepeat is increased.

**Figure 2.**
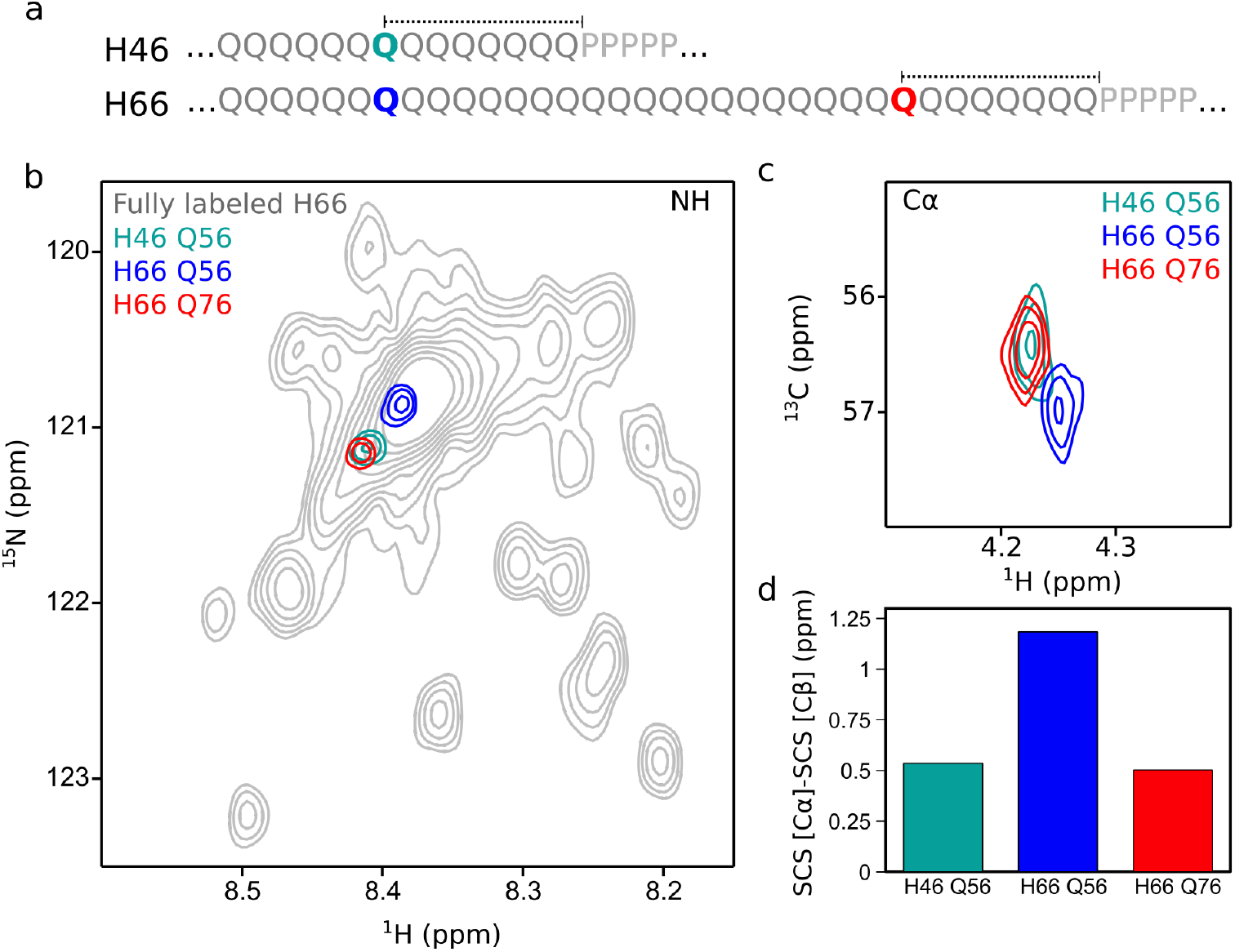
NMR analyses of H66 and comparison with H46. **(a)** Partial sequences of H46 and H66 indicating the positions of Q56 in H66 (green) and H46 (blue), and Q76 in H66 (red). This color code identifies the individual glutamines throughout the figure. Dashed lines highlight the equal distance of Q56 in H46 and Q76 in H66 to the PRR. Zoom of the ^15^N-HSQC (**b**) and ^13^C-HSQC (**c**) with individually superimposed colored SSIL spectra showing the poly-Q NH and Cα regions, respectively. (**d**) Secondary chemical shift (SCS) analysis of H66-Q56, H66-Q76 and H46-Q56, using experimental Cα and Cβ chemical shifts and a neighbor-corrected random-coil library.

**Figure 3.**
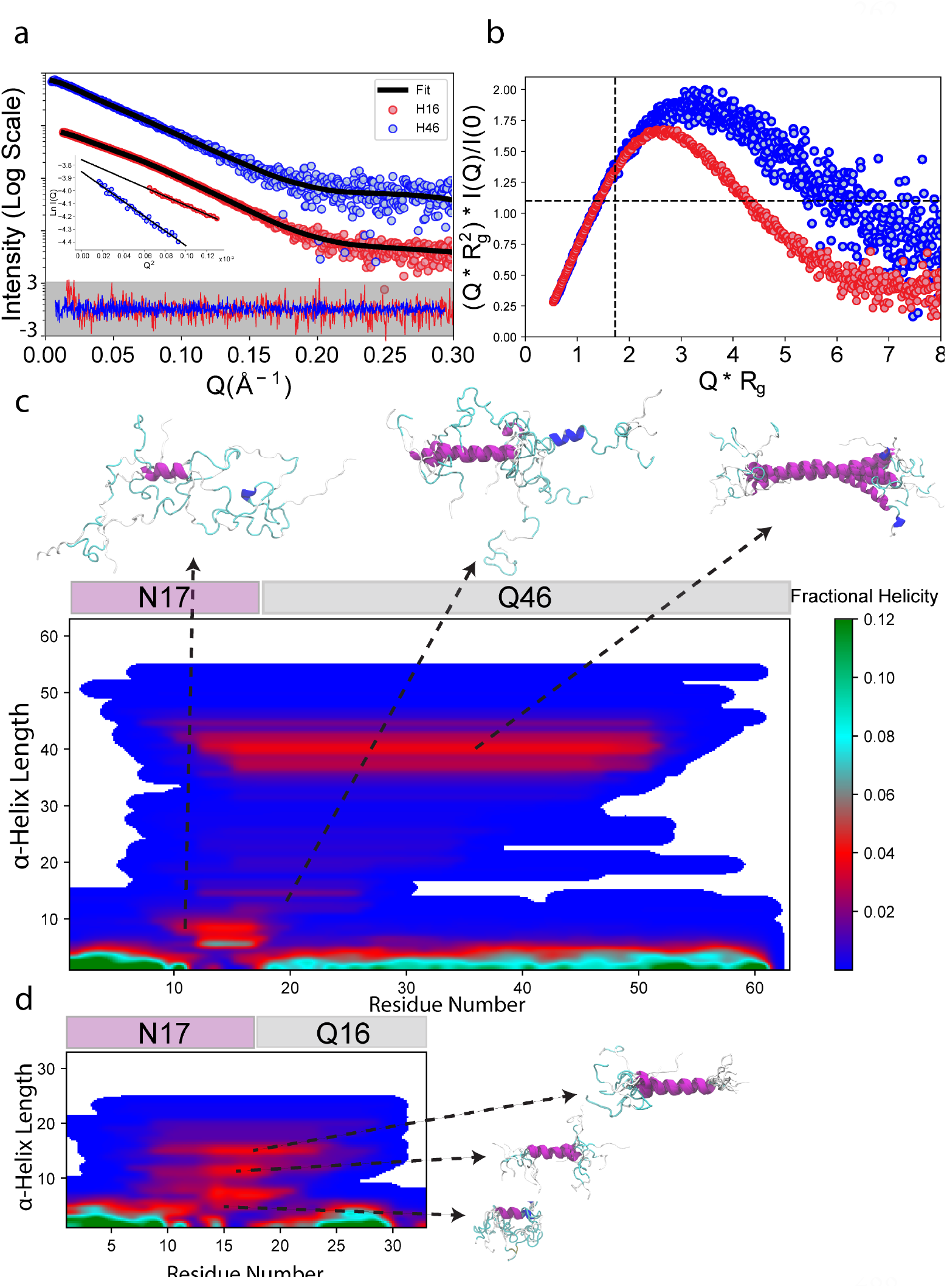
A structural model of pathogenic and non-pathogenic httex1 from the synergistic integration of NMR and SAXS data. **(a)** The SAXS intensity profiles for H16 (red) and H46 (blue) along with the theoretical profiles of EOM selected sub-ensembles (black lines). The residuals from EOM fitting are shown at the bottom. The inset shows the Guinier plots with linear fits as black lines. **(b)** The normalized Kratky plots for H16 (red) and H46 (blue) displaying a shift from the values expected for globular proteins (shown as black dashed lines) on both X and Y axes.**(c)** SS-maps calculated from the conformations selected during 100 cycles of EOM for H46 (top) and H16 (bottom). The population of the different α-helix lengths is shown according to the color code on the right. Some representative conformations with different lengths of helices are also shown.

### SAXS indicates that H46 is an elongated particle in solution with a large degree of flexibility

Small angle X-ray scattering (SAXS) was applied to derive the overall size of H46 in solution. To this end, we applied size-exclusion chromatography (SEC) coupled with SAXS, which allowed us to isolate the H46 peak from potential aggregates and partially digested sfGFP (Fig. S3). The analysis of the resulting profile (Fig. 3a and Table S2) indicates that H46 is a monomeric particle with a radius of gyration, *R*_*g*_, of 41.5 ± 0.5 Å according to Guinier analysis and a maximum dimension, *D*_*max*_, of 148 Å, calculated using pair-wise distance distribution, *p(r)*, analysis (Fig. S3c). Although the Kratky representation displayed a peak indicating the presence of a folded protein, which was attributed to the sfGFP, the intensity did not completely return to base line, suggesting that the httex1 part of H46 exhibited a high level of flexibility (Fig. 3b). The smooth decrease of the *p(r)* when approaching the *D*_*max*_ value and the departure in the maximum of the Kratky plot from the standard values of a globular protein substantiate the flexibility and the overall extendedness of the protein.

To further confirm that the overall extendedness is inherent to httex1, we performed an equivalent SAXS analysis for H16 (FigS. 3a, S3 and Table S2). Not surprisingly, the resulting SAXS profile of H16 indicated that the protein is a smaller particle (*R*_*g*_=32.9 ± 0.2 Å and *D*_*max*_ = 126 Å) in solution than H46 (Table S2). Importantly, H16 retained SAXS features corresponding to an extended and flexible particle observed for H46.

### H46 consists of a mixture of α-helical conformations of different lengths

The Cα chemical shifts and the SAXS data measured for H46 were integrated to derive a structural model of the protein. Similar to our previous study of the non-pathogenic H16, two families of ensembles were generated to capture the conformational influence of both flanking regions using a conformational sampling method for intrinsically disordered proteins (see Methods section for details)^38^. For the first family (N→C ensembles), starting with the ^10^AFESLKS^16^ region of N17 as partially structured, multiple ensembles of 5,000 conformations were built by successively including F17 and an increasing number of glutamines in the poly-Q tract (from Q18 to Q63) as partially structured, while the rest of the chain was considered to be fully disordered. An equivalent strategy was followed for the second family of ensembles (N←C ensembles) for which glutamines (from Q63 to Q18) were successively considered as partially structured starting from the poly-P tract. Note that using this building strategy, secondary structural propensities are naturally propagated due to the neighboring effects. For the resulting ensembles of each family and after building the side chains with the program SCWRL4^39^, averaged Cα chemical shifts were computed with SPARTA+^40^. For each family, the relative populations of these ensembles were optimized using the experimental Cα chemical shifts with Q55 as the boundary position (Fig. S4). Then, an NMR-compatible ensemble was built by randomly taking conformation from the selected ensembles with the appropriate populations (see Methods for details). This ensemble was further refined by integrating the SAXS data with the ensemble optimization method (EOM)^41,42^. For this, we included the crystallographic structure of sfGFP (PDBID: 3LVA) and the C-terminal His-tag to the individual conformations of the NMR-optimized ensemble to generate the pool of conformations used by EOM (see Methods). It must be noted that the H46 ensemble had an overall size (*R*_*g*_ = 42.7 Å) very similar to the experimentally determined one (*R*_*g*_, = 41.5 ± 0.5 Å), highlighting the quality of our starting model despite using local NMR information for the refinement. Sub-ensembles selected with EOM yielded an excellent fit to the experimental profile (χ^2^ = 0.2) (Fig. 3a). The resulting *R*_*g*_ distribution was quite broad and very similar to that of the initial ensemble, indicating that H46 is a highly flexible particle in solution and slightly more extended than the ensemble derived only using chemical shifts (Fig. S4).

The structural analysis of the NMR and SAXS compatible ensemble was performed with SS-map^43^, which displays the length, the residues involved and the population of the α-helices present in the ensemble in a comprehensible manner. This analysis substantiated the presence of a mixture of multiple helical conformations encompassing different sections of the H46 poly-Q tract (Fig. 3c). These α-helices were initiated in the last residues of the N17 domain (^14^LKSF^17^) and propagated along the tract. Interestingly, an enrichment of long α-helices encompassing around 40 residues and reaching up to Q52 was observed. The presence of these long stable helical conformations explains the steady plateau observed in the SCS analysis (Fig. 1d).

A similar structural refinement was performed for H16, using the previously reported NMR-refined ensemble^33^ and the SAXS data (Fig. 3a). The EOM fit yielded an excellent agreement with the experimental curve (χ^2^ = 1.12) and, as in the case of H46, the resulting sub-ensemble was slightly more elongated than that obtained using only the NMR information. We observed that H16 also consisted of a mixture of α-helical structures of different lengths, with prevalence for these encompassing a large fraction of the homorepeat (Fig. 3d).

### Side chain to backbone hydrogen bonds trigger and stabilize helical conformations in H46

It has been shown in non-pathogenic H16 and the androgen receptor that a network of hydrogen bonds involving backbone and side chains is at the origin of the helical propagation from the N-flanking region to the poly-Q tract^33,44^. Here, we investigated whether this effect is conserved in pathogenic httex1. To this end, we monitored the Cβ, Hβ, Cγ, Hγ and NHε chemical shifts, which could be unambiguously assigned through SSIL samples (Fig. 4a and S5). The two diastereotopic Hβ glutamine protons displayed resolved responses, with the difference in their chemical shifts increasing along the sequence. The maximum difference was observed for the last glutamines of the tract (Q62 and Q63) and those within the PRR (Q75 and Q91), while Q18 displayed the smallest one, featuring virtually degenerate Hβ chemical shifts (Fig. 4a, Fig. S5a). These observations indicate a correlation between the chemical shift difference and the level of disorder. Interestingly, Q21 exhibited three peaks that we attributed to the equilibrium between two conformations in slow exchange at the NMR timescale. One of the two conformations of Q21 displayed the spectroscopic features observed for Q18 (degenerate Hβ chemical shifts) and the other one those observed for the rest of the glutamines (distinct Hβ chemical shifts). This observation is in agreement with the two Cα peaks observed for Q21 (Fig. 1c), which were attributed to two conformations with different helical content.

**Figure 4.**
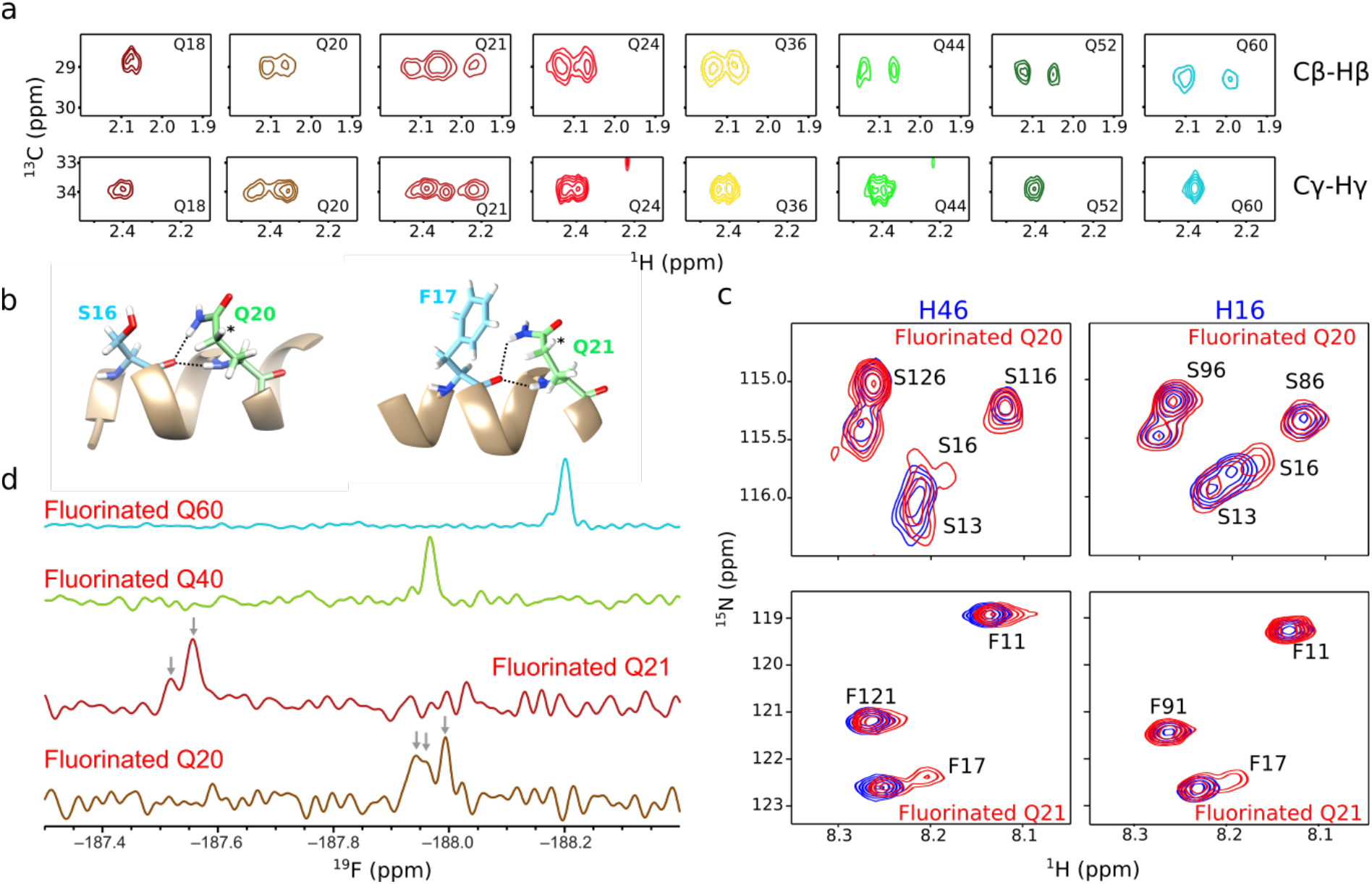
NMR analysis of H46 side chains. (**a**) Cβ-Hβ and Cγ-Hγ regions of the ^13^C-HSQC spectra of selected glutamines within the poly-Q. The spectra of Q60 display the standard behavior of disordered glutamines with non-degenerate and degenerate diastereotopic protons for Cβ-Hβ and Cγ-Hγ, respectively. (**b**) Structural model of the N17-poly-Q coupling showing bifurcated H-bonds between S16 and Q20 (left) and F17 and Q21 (right). The asterisk indicates the proton substituted with a fluorine atom when 4F-Gln was incorporated. (**c**) Zoom of ^1^H-^15^N HSQC spectra of H46 (left panels) and H16 (right panels) samples labeled with either ^15^N-Ser or ^15^N-Phe. In red, spectra of samples with fluorinated glutamine at position Q20 (upper panels) or Q21 (lower panels). Non-fluorinated samples are colored in blue. (**d**) 1D ^19^F spectra of H46 samples with fluorinated glutamines at positions Q20, Q21, Q40 or Q60.

The Hγ signals followed an inverse trend than that observed for Hβ. The last glutamines of the tract and those located in the PRR presented a single correlation, corresponding to degenerate diastereotopic Hγ chemical shifts, as expected for a flexible glutamine side chain (Fig. 4a and S5b). Conversely, the majority of glutamines of the homorepeat (up to Q48) displayed a small but measurable difference between the Hγ chemical shifts, suggesting a transient rigidification of the side chain. Again, in line with the Hβ signals, Q18, Q20 and Q21 were exceptions to this behavior. While Q18 exhibited a single Cγ-Hγ peak in the ^13^C-HSQC, Q21 displayed three peaks, substantiating the equilibrium between the two previously eluded conformational states. Although Q20 only exhibited two Cγ-Hγ peaks, their difference in intensity suggested a similar situation as for Q21, but with the Cγ-Hγ peak of the conformational state with degenerate Hγ chemical shifts overlapping with one of the Cγ-Hγ peaks of the other state.

These results confirm previous observations for a non-pathological version of httex1 and the androgen receptor^33,44^, demonstrating the presence of *i*→*i*+4 bifurcated hydrogen bonds structurally connecting the first residues of the poly-Q tract with the upstream flanking region (Fig. 4b). Furthermore, our data indicate that these hydrogen bonds are present, although to a lower extent, along the homorepeat, incorporating an additional mechanism for structural stabilization.

### 2S,4R-Fluoroglutamine (4F-Gln) as a new probe to study the structure and dynamics of httex1

In order to confirm the different conformational behavior of glutamine side chains along the poly-Q tract and benefiting from the SSIL methodology, a fluorinated glutamine was site-specifically incorporated at different positions of H46 (Fig. 1a). Consequently, we synthesized with high yield and stereospecificity 2S,4R-fluoroglutamine (4F-Gln), in which a fluorine atom replaced a hydrogen atom on Cγ (Fig. S6)^45^. This 4F-Gln was successfully loaded onto the tRNA_CUA_ using the yeast glutaminyl-tRNA synthetase with similar yields as for canonical glutamine (Fig. S7), suggesting that the fluorine atom is not changing the structural and electronic properties of glutamine, enabling the recognition by the enzyme^31^.

4F-Gln was used in two sets of experiments. First, we used 4F-Gln to substantiate the participation of Q20 and Q21 in the hydrogen bond network that propagates helicity in the homorepeat. To this end, we incorporated 4F-Gln in positions Q20 or Q21 in two H46 samples that were also isotopically labeled with ^15^N-Ser or ^15^N-Phe, respectively (Fig. 4c left panels and Fig. S8). When comparing their ^15^N-HSQC spectra with those of non-fluorinated H46 (Fig. 4c, left panels), substantial changes could be observed in S16 and F17 upon Q20 or Q21 fluorination, respectively. The presence of a fluorinated glutamine in position 20 produced a slight chemical shift change of S16, while fluorination of Q21 induced a stronger effect on F17, leading to the appearance of a second peak. Furthermore, no chemical shift changes were observed for the other serines and phenylalanines of H46, highlighting the specificity of the interaction. Importantly, similar observations were made when 4F-Gln was incorporated in the same positions in H16, in samples simultaneously labeled with ^15^N-Ser and ^15^N-Phe (Fig. 4c right panels and Fig. S8). Indeed, when Q20 was fluorinated in the non-pathogenic form, not only S16 was affected, but F17 was also perturbed (Fig. S8). Fluorination of Q21 again resulted in the appearance of a second F17 signal and slightly affected S16. These data underline the structural coupling between Q20 and S16 as well as Q21 and F17, in both pathogenic and non-pathogenic forms of httex1.

In a second set of experiments, we incorporated 4F-Gln in four H46 positions strategically located at the beginning (Q20 and Q21), the middle (Q40) and the end (Q60) of the poly-Q tract to be monitored by 1D ^19^F-NMR (Fig. 1a). Note that the ^19^F chemical shift is exquisitely sensitive to small differences in the electronic surroundings of the fluorine nucleus, and thus an excellent reporter on biomolecular structure and dynamics^46,47^. Strong differences in ^19^F chemical shifts were observed for the four positions despite the homogeneity of the amino acid sequence (Fig. 4b). This demonstrates important structural changes along the poly-Q tract, most likely linked to the amount of helical content. The chemical shift of Q21 was particularly high, most probably due to its proximity to the ring currents exerted by F17 in the hydrogen-bonded form (Fig 4b and 4d).

Interestingly, multiple ^19^F-NMR resonances were observed for Q20 and Q21. For Q20, three distinct responses were measured, including two signals with very close chemical shifts that could only just be resolved. In line with the Cγ-Hγ peaks, it can be concluded that the Q20 side chain adopts at least two different conformations in a slow exchange regime. Given the sensitivity of the ^19^F chemical shift, the two closely resonating signals can be explained by the fluorine sensing the different conformational states of neighboring glutamines. For Q21, a weak second ^19^F signal could be detected, in agreement with the two populations previously identified in the Cα-Hα, Cβ-Hβ and Cγ-Hγ correlations. Altogether, the site-specific incorporation of 4F-Gln substantiates the structural link of N17 with the poly-Q tract, the distinct glutamine side-chain conformational preference along the poly-Q tract, and the presence of multiple conformations in the homorepeat.

### Molecular dynamics simulations provides insights into the helix propagation and stabilization mechanisms in httex1

We performed Gaussian accelerated molecular dynamics (GaMD)^48^ simulations to understand the secondary structure propensities of httex1 and get further insights into the role of N17 in inducing helicity. Previous MD studies of httex1 have yielded vastly different results ranging from completely α-helical to highly β-strand rich structures, most likely due to the use of non-adapted force-fields^49^. To simulate a httex1 fragment encompassing N17, the polyQ tract and five prolines, we used the recently developed ff99SBws-STQ force-field, which has been refined to describe the secondary structure propensities of low-complexity proteins^50^. In all the 8 independent MD simulations, with an aggregated time of ≈20 µs, we observed that the poly-Q tract sampled a wide conformational landscape and mainly adopted α-helical and disordered conformations, while N17 presented a higher helical propensity (Fig. 5a and S9a). Interestingly, several helical folding and melting events could be observed along these trajectories. For some of the simulations, we observed the formation of long α-helices that spanned almost the whole length of N17 and poly-Q tract and were only absent in the residues close to the poly-P region (Fig. 5a). In line with the directionality observed experimentally, the process of α-helix melting systematically occurred from C to N, as observed in the 1600-1800 ns period in Fig. 5a, while α-helices in httex1 preferentially grew from N to C (Fig. S10). The spontaneous formation of short α-helical conformations unconnected with N17 was also observed along the trajectories (*e*.*g*., frames 2000-2300 in Fig. 5a), although they dissolved relatively fast. This suggests a small inherent α-helical propensity of poly-Q tracts.

**Figure 5.**
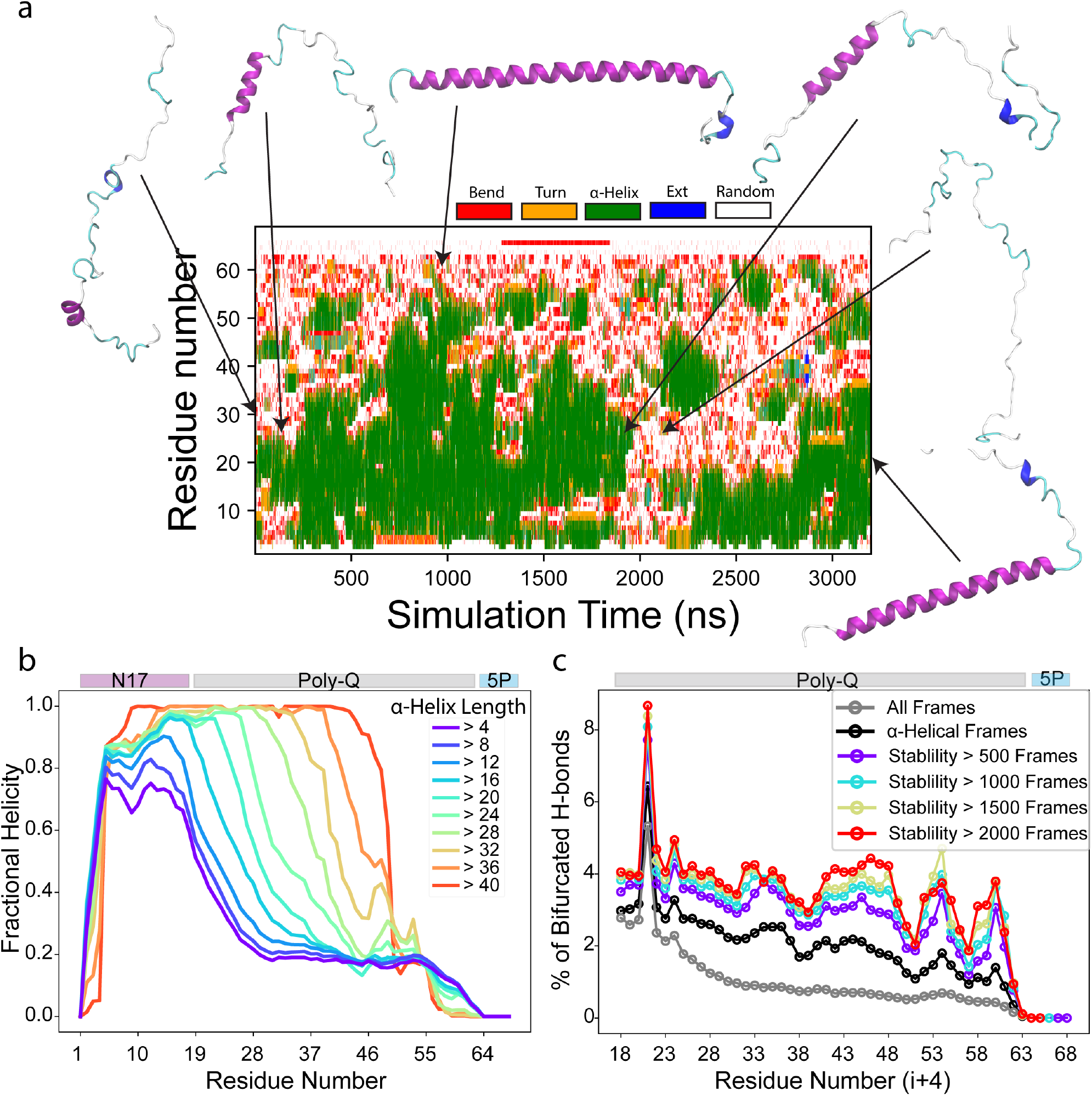
Insights into the conformational landscape of httex1 with MD simulations. **(a)** The per-residue secondary structure plot for one of the GaMD trajectories showing a transition from an almost completely random-coil conformation to a long α-helix and back to random coil. **(b)** The reweighted fractional helicity calculated using frames with increasing minimum α-helix length ranging from 4 to 40. **(c)** The percentage of frames with bifurcated hydrogen bonds for each residue of the poly-Q tract using all frames (gray), only the frames where the fragment (*i →i+4*) was in helical conformation (black) and the segments of the trajectory where the fragment (*i →i+4*) formed a stable α-helix for an increasing number of frames ranging from 500 to 2000.

The weighted average per-residue fractional helicity of all the simulations indicated that the poly-Q tract adopted higher α-helical propensity when closer to N17 (Fig. S9b). Moving further from N17, the helicity decreased until Q30 from where it remained flat until Q55 to finally decrease when approaching the poly-P. Importantly, this behavior was qualitatively very similar to the experimentally determined α-helix propensity of httex1 (Fig. 1d and 1e). This suggests that our simulations captured the structural mechanisms present in httex1, although the relative population of α-helix in the poly-Q tract was underestimated. When analyzing the frames with α-helices with increasing length (from 4 to 40), we observed a gradual increase in fractional helicity from N- to C-terminus, substantiating the structural propagation from N17 towards the poly-Q (Fig. 5b).

We analyzed the trajectories for the existence of *i→i+4* bifurcated hydrogen bonds. Interestingly, they were found throughout the poly-Q, although they were more abundant in the beginning of the tract where the hydrogen bonds were formed with the residues of N17 region and their number slowly decreased along the tract (Fig. 5c). Residue Q21 presented the highest propensity to form bifurcated hydrogen bonds with F17. Taken together, the trend of bifurcated hydrogen bonds agrees with the results of the appearance of doublets in Cγ-Hγ correlations and the results from specifically fluorinated glutamines probing the initial residues of the poly-Q (Fig. 4). Then, we analyzed the correlation between bifurcate hydrogen bonds and α-helix stability. Not surprisingly, the percentage of these hydrogen bonds was higher in frames where the segment (*i →i+4*) adopted a helical conformation. Importantly, the population consistently increased with the stability of the helix, suggesting that bifurcated hydrogen bonds stabilize α-helical conformations in httex1 (Fig. 5c).

### The structure of the poly-Q governs aggregation kinetics and fibril structure of httex1

The hydrogen bond network connecting N17 and the poly-Q tract enabled the design of point mutants in H16 altering the level of structure in the homorepeat while preserving its length^33^. Its application to H46 allowed the interrogation of the relative relevance of the helical content and the poly-Q tract length for the aggregation propensity in a pathogenic variant of httex1. With this aim, two N17 mutants of H46 were produced (Fig. 1a). First, by substituting ^16^SF^17^ by ^16^GG^17^ (LKGG-H46), we hamper the H-bond network connecting both domains. Second, when ^15^KS^16^ were replaced by ^15^LL^16^ (LLLF-H46), the hydrogen bond network was strengthened by incorporating two additional large hydrophobic residues. Uniformly labeled and SSIL samples of these mutants were produced and analyzed by NMR. On one hand, LKGG-H46 glutamine ^15^N-HSQC signals collapsed in a broad, high-intensity, downfield-shifted peak, proving a substantial loss of helicity in comparison with the wild-type H46 (Fig. 6a). Interestingly, this broad peak did not overlap with the positions corresponding to fully unstructured glutamines, which were shifted further downfield. This indicates that poly-Q, even when disconnected from the flanking region, contains a small intrinsic propensity for helical conformations, in agreement with our MD simulations and previous studies^35^. On the other hand, the ^15^N-HSQC spectrum of fully labeled LLLF-H46 displayed a more dispersed density of glutamine peaks and an additional upfield density, pointing to a helicity increase of the poly-Q tract (Fig. 6b). The detailed analysis of NH, NεH_2_ and Cα signals of Q18, Q20 and Q21 SSIL samples from both mutants confirmed the decrease in structuration of LKGG-H46 and the increase in helicity in LLLF-H46, in comparison with the wild-type form (H46) (Fig 6a,b and Fig. S11). Importantly, Q21 in LLLF-H46 displayed signatures of two conformations as also observed for the wild-type. Interestingly, this residue exhibited two NHε correlation peaks, suggesting the formation of a stronger bifurcated hydrogen bond than in the wild-type. Unfortunately, a single Cα peak, corresponding to the less helical conformation, was observed for Q21 in this mutant, suggesting an unfavorable exchange regime for the NMR detection.

**Figure 6.**
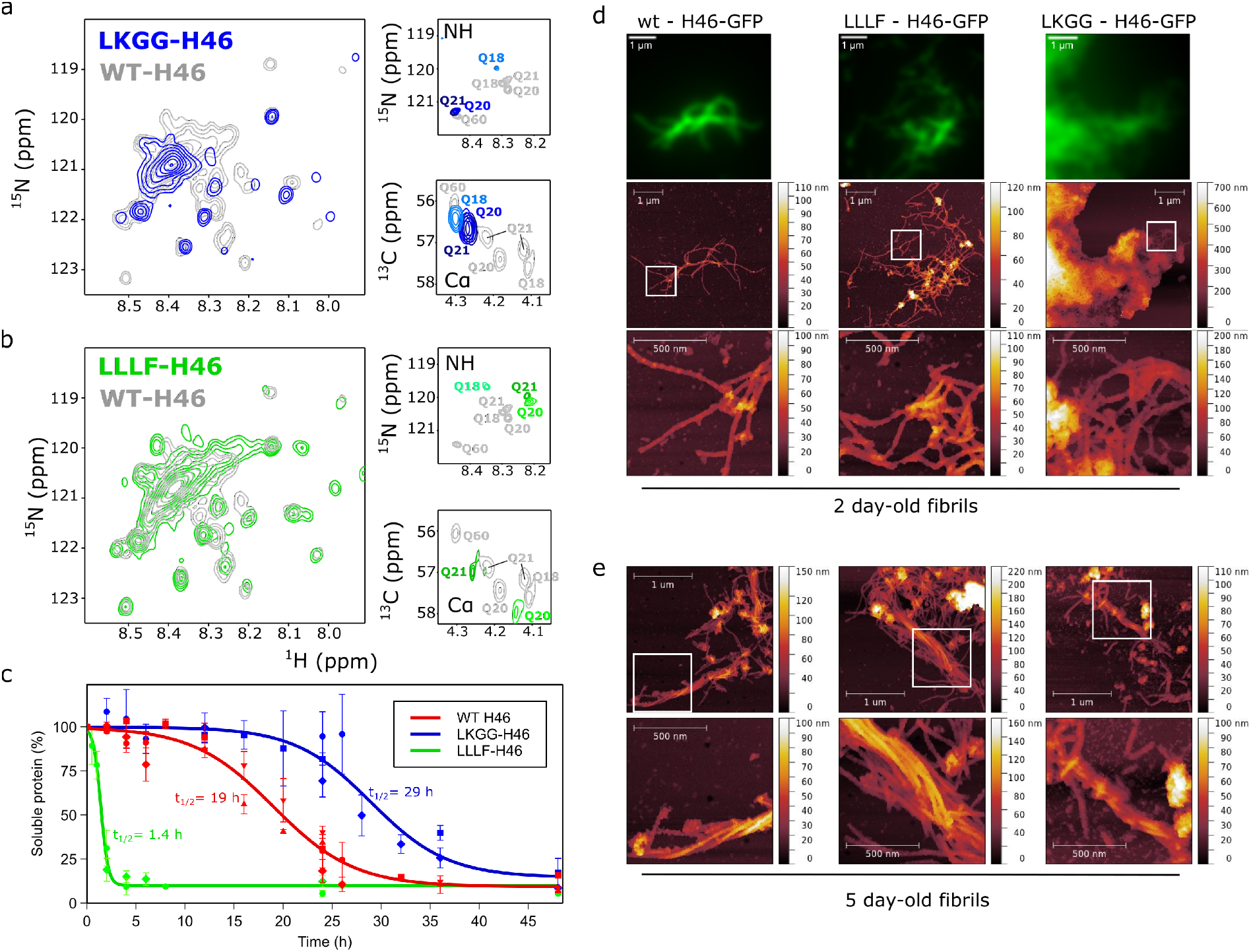
Structural and fibrillation analyses of H46 N17 mutants. **(a, b)** Overlay of the glutamine region of the ^15^N-HSQC of fully labeled H46 (gray) with the N17 mutants LKGG-H46 (blue) and LLLF-H46 (green), as well as zooms of the NH and Cα regions of the corresponding Q18, Q20 and Q21 SSIL samples. (**c**) Time course of aggregation for wild-type, LKGG- and LLLF-H46 (15 μM) at 37 °C. Each data point corresponds to the mean and associated standard deviation calculated from three replicates. Symbols represent different independent experiments. Half-time (t_1/2_) values calculated for each H46 species are indicated according to the color code shown in the legend. (**d**) Fluorescence microscopy (upper panels) and AFM (middle and lower panels) images of 2-day-old fibrils of wild-type, LLLF- and LKGG-H46. Each fluorescence image corresponds to the average of 150 pictures. (**e**) AFM images of 5-day-old fibrils of three H46 species. White squares indicate the zoom region displayed in the panels below.

Once the structural features of the H46 mutants were confirmed, their propensities to aggregate were measured. For this, 15 µM samples of H46, LKGG-H46 and LLLF-H46 were incubated at 37°C and the soluble fractions of the proteins were analyzed by SDS-PAGE over a period of 48 h (Fig. 6c and Fig. S12). Several essays were performed to better sample the monitored period. H46 presented a moderate aggregation propensity, with a systematic decrease of the soluble fraction from the first hours of incubation that almost disappeared after 48 h. When the soluble fraction was fitted to a reverse sigmoid function an aggregation half-time, t_1/2_, of 19 h was obtained. A much stronger aggregation propensity was observed for the LLLF-H46 mutant, for which a t_1/2_ of only 1.4 h was derived. These observations indicate that increasing the α-helical stability strongly enhances the aggregation propensity of httex1. When the same experiment was performed with the LKGG-H46 mutant, which has a more disordered poly-Q tract similar to pure poly-Q peptides^51^, the first signs of aggregation occur after 20 h of incubation and a t_1/2_ of 29 h was obtained. The presence of sfGFP fused to the C-terminus of the httex1 variants, which certainly slows down aggregation, makes the quantitative comparison with previous studies difficult^52^. However, the relative aggregation propensity of the three httex1 forms can be unambiguously obtained.

The morphology of the aggregates for H46 and the two mutants after 48 and 120 h were investigated by correlative atomic force microscopy (AFM) – total internal reflection fluorescence (TIRF) (Fig. 6d,e and S13a,b). The intrinsic fluorescence of the proteins allowed the easier localization of the aggregates on the silica surface and indicated that no proteolytic activity preceded the aggregation process of the samples. The inspection of H46 micrographs measured after 48 h of incubation revealed the presence of typical amyloid structures with interconnected fibrils, often presenting a length larger than 1 μm (Fig. 6d, upper panels, left row). Those fibrils exhibited a heterogeneous morphology, with notable variations in width and height, similarly to those recently described^53,54^. Interestingly, LLLF-H46 aggregates displayed similar features, although the presence of fibrils was substantially more abundant (Fig. 6d, middle panels). Indeed, large aggregates containing long fibrils were easily found in the sample (Fig S13a). Again, width variations were often observed along the fibrils. The analysis of LKGG-H46 preparation revealed a different behavior. Considerably larger and more heterogeneous aggregates, presenting often well-defined limits, were observed for this mutant. Images indicated the presence of very short fibrils, although relatively long isolated fibrils with a similar morphology than the wild-type and the LLLF-H46 were found in the boundaries of the heterogeneous aggregates.

To verify whether the morphology was maintained after longer maturation times, 5-day-old fibrils were also imaged (Fig. 6e and Fig. S13b). Long fibrils were found in wild-type and LLLF-H46 preparations and, interestingly, elongated bundle structures involving several paired filaments were observed. However, these coiled structures were seldom found in LKGG-H46 aggregates and, when observed, they were less ordered (Fig. 6e). Indeed, fibrils around LKGG-H46 bundles were fragmented and small aggregates could be detected along the whole sample, suggesting reduced fibril stability. Altogether, aggregation and AFM experiments demonstrate that the structural properties of httex1 exert a stronger influence on the aggregation propensity and the final form of the fibrils than the poly-Q tract length.

## Discussion

Huntington’s disease is the most notorious example of the poly-Q-related diseases and findings connecting structure and disease unveiled for huntingtin are most probably applicable to the whole family of pathologies^1^. In this study, we have characterized at high resolution a pathogenic form of httex1 containing 46 consecutive glutamines, a system that is out of the reach for traditional structural biology approaches. We demonstrate that SSIL can be systematically applied to highly repetitive sequences, such as H66, and the only limitations arise from the capacity to purify the protein and preserve its stability in solution.

The systematic application of SSIL to H46 demonstrates that this protein retains the α-helical conformation previously observed for non-pathogenic versions of httex1^26,29,33^. Using circular dichroism, α-helical propensity in pathogenic httex1 had been previously identified for httex1 with up to 55 glutamines^27^. Moreover, it was shown that the helical content increased concomitantly with the length of the tract. However, the low-resolution of circular dichroism hampered the analysis of the extent and stability of helical conformations. One of the most striking observations of our study is the fairly flat plateau of measured secondary chemical shifts along a large fraction of the poly-Q tract (Fig. 1d). These chemical shifts are consistent with the coexistence of multiple partially formed α-helices of different lengths, spanning almost the complete poly-Q tract. However, these helices are not equally populated and long α-helices are prevalent according to our synergistic analysis of NMR and SAXS data (Fig. 3). This behavior evidences that the helices, which are formed through a nucleation process triggered by the interaction between the N17 and the first glutamines of the tract, are cooperatively propagated along the homorepeat. A faster decrease in the SCS values would be expected if all helical lengths would display a similar thermodynamic stability. The comparison of the NMR observables for H16 with those of H46 also substantiates the helical stabilization with the poly-Q length. For instance, despite both proteins having the same sequence context, NH, Cα and NεH_2_ chemical shifts for initial glutamines of the tract are systematically shifted towards more helical conformations in H46 than in H16 (Fig. S1). Moreover, experiments performed on H66 demonstrate that α-helical conformations are maintained for long poly-Q tracts and the distance of individual glutamines to the PRR defines their helical propensity (Fig. 2).

From a structural perspective, the α-helical stability found in httex1 could arise from the formation of bifurcate hydrogen bonds all along the homorepeat. Indeed, up to Q48, the diastereotopic Hγ protons have non-degenerate chemical shifts, although the chemical shift difference decreases along the repeat (Fig. S5). An opposite trend is observed for the diastereotopic Hβ protons, which present increasing chemical shift differences when reaching the less structured glutamines of H46. These spectroscopic features have been associated with the formation of bifurcated hydrogen bonds and the concomitant rigidification of glutamine side chains^33,44^. Our MD simulations indicated that the percentage of bifurcated hydrogen bonds is correlated with the stability of α-helical conformations. The ensemble of these observations suggests that glutamine side chains are actively involved in the stabilization of the helical conformation of the poly-Q tract and that the strength of this mechanism declines when approaching the PRR.

Partial structuration in the poly-Q implies that individual glutamines co-exist in (at least) two different conformational states. A similar conclusion was reached by EPR experiments after incorporating stable radicals in several positions of a pathogenic httex1^27^. In that study, two dynamic regimes were identified in N17 and poly-Q tract residues, but not in glutamines in the PRR. However, structural details of this conformational fluctuation could not be unveiled. Several NMR signatures collected for H46 using SSIL samples, which can be rationalized through the MD simulations, define the structural bases of this conformational equilibrium. These include, among others, the two Cα-Hα peaks for Q21 (Fig. 1c), the two sets of Cβ-Hβ and Cγ-Hγ peaks for Q20 and Q21 (Fig. 4a), and the multiple ^19^F-NMR frequencies also detected for these two residues. Remarkably, the co-existence of a conformational equilibrium is also manifested in most of the other glutamines of the tract, which exhibit the previously eluded non-degenerate Cγ-Hγ peaks. Our observations demonstrate that httex1 fluctuates between a rigid α-helical conformation, which is stabilized with bifurcate hydrogen bonds, and a more disordered state. Furthermore, our data suggest that the dynamic regime also changes in conjunction with the stability of the helical conformation. While the first glutamines of the tract, which have the highest helical propensity, exhibit a slow exchange on the NMR frequency timescale, a fast exchange regime is observed for the other helical glutamines. This asymmetric behavior suggests that helix unwinding is initiated in the proximity of the PRR and progresses towards the N-terminus, substantiating the previously proposed protective role exerted by the PRR^34,35,55,56^.

The synergistic combination of NMR and SAXS data has enabled the elucidation of an ensemble model of H46^57^, showing that the presence of long α-helices determines the overall shape of httex1. The refined ensemble indicates that H46 is a flexible elongated particle in solution and that the overall size is correlated with the length of the poly-Q tract (Fig. 3c). Importantly, our structural model is in contradiction to previously reported compact models of httex1^25,26,49,58^. In these models, compactness is driven by extensive fuzzy contacts between N17 and the poly-Q, a situation that is not compatible with our data. Despite the overall extendedness of the httex1 structure derived here, our ensemble description requires that a large fraction of the protein ensemble remains disordered. This disorder explains the fluorescence transfer efficiency observed in smFRET measurements^25^ and the lack of permanent hydrogen bonds observed in NMR hydrogen deuterium exchange (HDX) experiments^26^.

The similarity of the mechanisms defining the structure of non-pathogenic and pathogenic forms of httex1 validates ‘linear lattice’ as the model accounting for the existence of the pathological threshold^20,22–24^. According to this model, the expansion of the poly-Q induces a systematic increase of the toxicity of httex1 and, beyond the pathological threshold, triggers cytotoxicity and neuronal aggregation. Importantly, our study provides a structural perspective for this model. Our results demonstrate that the poly-Q extension is associated with an increase in the length and stability of the coexisting helical conformations. This is relevant as our aggregation experiments unambiguously show that the α-helical content and not the homorepeat length is the key factor promoting aggregation. Although N17 has been demonstrated to be the aggregation-triggering domain^18,30,34^, httex1 becomes aggregation-prone when N17 is structurally coupled to the poly-Q tract. This suggests that when structurally uncoupled, such as in the LKGG-H46 mutant, partially helical N17 is still able to oligomerize, but the resulting oligomers are less stable and aggregation propensity is notably reduced. Conversely, strengthening the structural coupling between both domains stabilizes the helical content of the protein and accelerates aggregation, as observed for LLLF-H46. In line with these observations, previous aggregation experiments with httex1 analogues modifying the poly-Q tract structure highlighted the relevance of the homorepeat conformational preferences in defining the aggregation propensity^15,59^. In addition to modified aggregation kinetics, we have observed that mutations in the flanking regions also result in fibers with distinct morphologies, with the LKGG-H46 mutant forming shorter amyloids and exhibiting less capacity to associate to form bundles. Polymorphism in httex1 aggregates has been previously observed *in vivo* and *in vitro* when modifying the experimental conditions or when deleting flanking regions^15,52,60–62^. This polymorphism has been associated to the contribution of N17 and the PRR to the fibril packaging^62,63^. Conversely, the amyloid core is consistently formed by antiparallel poly-Q stretches forming β-sheet monolayers connected through interdigitated glutamine side chains^63,64^.

In light of our observations, we can speculate about the structural bases of the mechanisms leading to the pathology (Fig. 7). The helical propensity in the poly-Q tract may facilitate intermolecular assemblies through coiled coil (CC) interactions. Indeed, more than 60% of the described httex1 partners have, or are predicted to have, CCs^65^. Furthermore, CC formation has also been suggested to be an important step for the oligomerization and subsequent aggregation of httex1^17,19^. In the non-pathological scenario, the short length and relatively low stability of the poly-Q helical conformations precisely define the selectivity of partners recognized by httex1. Moreover, resulting httex1 oligomers exhibit reduced stability. When the number of glutamines exceeds the pathological threshold, there is a concomitant population increase of long α-helical conformations. These long poly-Q helices could still interact with their physiological partners, although most probably with different thermodynamic properties compared to the non-pathological scenario. However, they could also associate with other non-physiological partners, sequester them and perturb crucial signaling and metabolic pathways, inducing the symptoms associated to HD^2^. In terms of the fibrillation capacity, longer helices can form more stable oligomers though CC interactions. These oligomers can eventually nucleate the formation of the amyloidogenic fibrils^17,66^ or being the source of cytotoxicity by sequestering crucial cellular components^67^.

**Figure 7.**
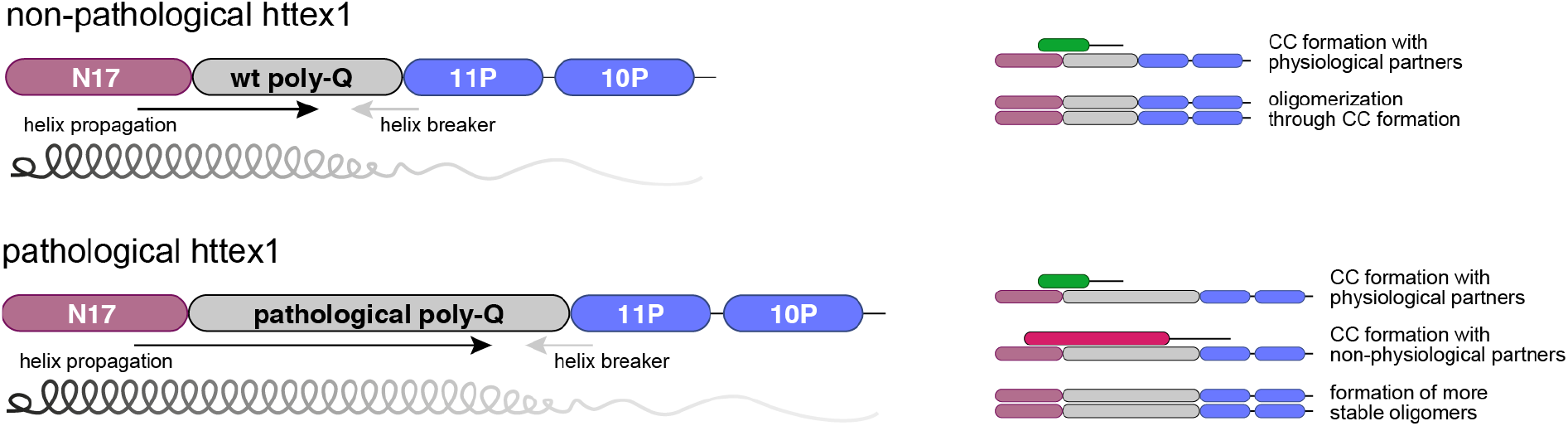
Scheme illustrating the structural influences within non-pathogenic and pathogenic httex1 and their respective modes of interaction. (**Top, non-pathogenic httex1**) The poly-Q tract of non-pathogenic httex1 experiences opposing structural effects from the N17 (α-helix propagation; black arrow) and the PRR (helix breaking; gray arrow). The gradually shaded helix represents the decrease in helical propensity. Due to its helical conformation, non-pathogenic httex1 can form coiled coil (CCs) with physiological partners or with other httex1 molecules, resulting in oligomers. (**Bottom, pathogenic httex1**) Pathogenic httex1 experiences the same structural effects, however, helix propagation outweighs the helix-breaking effect coming from the PRR and enhances the formation of CCs. Pathogenic httex1 can still interact with its physiological partners *via* CC formation, but it can also interact with non-physiological partners and form more stable oligomers, which eventually drive to fibrils.

Altogether, this study shows that the expansion of the poly-Q tract in pathogenic httex1 is associated with an increase in the length and stability of α-helical conformations, which are the main driving force for the enhanced aggregation propensity. Our results provide a high-resolution structural perspective of the pathological threshold in HD that goes beyond the length of the poly-Q tract by underlining the associated conformational preferences. The generalization of these observations to the other poly-Q related diseases remains to be unveiled. However, the possibility to explore their associated proteins at the residue level and independently of the length of the homorepeat paves the way to a detailed structural understanding of the origin of these pathologies.

## Materials and Methods

### Huntingtin exon1 constructs

All plasmids were prepared as previously described^33^. Synthetic genes of wild-type huntingtin exon1 with 16, 46 and 66 consecutive glutamines (H16, H46 and H66 respectively) or H46 and H66 carrying the amber codon (TAG) instead of the glutamine codon, *e*.*g*. Q18 (H46Q18), were ordered from GeneArt. Following this strategy, 16 amber mutants for H46 and two for H66 were purchased: Q18, Q20, Q21, etc. Synthetic genes of the structural H46 mutants (LKGG-H46 and LLLF-H46) and their corresponding amber codon mutants (Q18, Q20, Q21) were also ordered from GeneArt. All genes were cloned into pIVEX 2.3d, giving rise to pIVEX-httex1-3C-sfGFP-His_6_ and mutants. The sequence of all plasmids was confirmed by sequencing by GENEWIZ.

### Synthesis of 2S,4R-fluoroglutamine

The synthesis of the 2S,4R-fluoroglutamine ((2S, 4R)-2,5-Diamino-4-fluoro-5-oxopentanoic acid), 4F-Gln, was performed as detailed by Qu *et al*.^45^. The purity and the enantiomeric excess (98%) were evaluated by ^1^H- and ^19^F-NMR (see Fig. S6 in Supplementary information).

### Preparation and aminoacylation of suppressor tRNA_CUA_

A tRNA_CUA_/tRNA synthetase pair based on the Gln2 tRNA and glutamine ligase GLN4 from *Saccharomyces cerevisiae* was prepared in house as previously described^32^. Briefly, the artificial suppressor tRNA_CUA_ was transcribed *in vitro* and purified by phenol-chloroform extraction. Prior to use, the suppressor tRNA_CUA_ was refolded in 100 mM HEPES-KOH pH 7.5, 10 mM KCl at 70°C for 5 min and a final concentration of 5 mM MgCl_2_ was added just before the reaction was placed on ice. The refolded tRNA_CUA_ was then aminoacylated with [^15^N,^13^C]-glutamine (CortecNet) in a standard aminoacylation reaction: 20 µM tRNA_CUA_, 0.5 µM GLN4, 0.1 mM [^15^N, ^13^C]-Gln (or 0.5 mM of 4F-Gln) in 100 mM HEPES-KOH pH 7.5, 10 mM KCl, 20 mM MgCl_2_, 1 mM DTT and 10 mM ATP^37^. After incubation at 37°C for 1 hour GLN4 was removed by addition of glutathione beads and loaded suppressor tRNA_CUA_ was precipitated with 300 mM sodium acetate pH 5.2 and 2.5 volumes of 96% EtOH at −80°C and stored as dry pellets at −20°C. Successful loading was confirmed by urea-PAGE (6.5% acrylamide 19:1, 8 M urea, 100 mM sodium acetate pH 5.2).

### Standard cell-free expression conditions

Lysate was prepared as previously described^33^ and based on the *Escherichia coli* strain BL21 Star (DE3)::RF1-CBD_3_, a gift from Gottfried Otting (Australian National University, Canberra, Australia)^68^. Cell-free protein expression was performed in batch mode as described by Apponyi *et al*.^69^. The standard reaction mixture consisted of the following components: 55 mM HEPES-KOH (pH 7.5), 1.2 mM ATP, 0.8 mM each of CTP, GTP and UTP, 1.7 mM DTT, 0.175 mg/mL *E. coli* total tRNA mixture (from strain MRE600), 0.64 mM cAMP, 27.5 mM ammonium acetate, 68 µM 1-5-formyl-5,6,7,8-tetrahydrofolic acid (folinic acid), 1 mM of each of the 20 amino acids, 80 mM creatine phosphate (CP), 250 µg/mL creatine kinase (CK), plasmid (16 µg/mL) and 22.5% (v/v) S30 extract. The concentrations of magnesium acetate (5-20 mM) and potassium glutamate (60-200 mM) were adjusted for each new batch of S30 extract. A titration of both compounds was performed to obtain the maximum yield.

### Preparation of NMR samples

Samples for NMR studies were produced in cell-free at 5-15 mL scale and incubated at 23°C and 450 rpm in a thermomixer for 4 h. Uniformly labeled NMR samples were obtained by substituting the standard amino acid mix with 3 mg/mL [^15^N, ^13^C]-labeled ISOGRO^40^ (an algal extract lacking four amino acids: Asn, Cys, Gln and Trp) and additionally supplying [^15^N, ^13^C]-labeled Asn, Cys and Trp (1 mM each) and 4 mM Gln. H46 samples in which only certain amino acids were selectively labeled (Ala and Lys; Gly, Ser and Arg; Leu and Glu; and Phe) were prepared by substituting the respective amino acids for the [^15^N, ^13^C]-labeled ones. To enable the labeling of glutamates, potassium glutamate was substituted with potassium acetate, which was optimized by testing a range of concentrations^36^. To produce site-specifically labeled samples, the standard reaction mixture was slightly modified. Instead of adding 1 mM of each amino acid, proline and glutamine were substituted by deuterated versions (Eurisotop) and used at 2 or 4 mM, respectively. 10 µM of [^15^N, ^13^C]-Gln or 4F-Gln suppressor tRNA_CUA_ were added for suppressed samples. The same procedures were used for the preparation of H66 samples.

### Expression of H16 in *E. coli*

*Escherichia coli* BL21 (DE3) transformed with H16 construct was grown in LB medium supplemented with 50 μg/mL kanamycin at 37ºC under stirring. When an OD_600nm_ 0.7 was reached, the culture was induced using 1 mM IPTG and grown for 24 hours at 23ºC. The cell pellet was collected by centrifugation at 5,000 x g for 15 minutes at 4ºC and resuspended in 10 mL buffer A (50 mM Tris, 1000 mM NaCl, pH 8.5), supplemented with cOmplete EDTA free protease inhibitor tablet (Roche), per 1 L of expression volume. Cells were lysed by sonication at 35% for 2 minutes with on-off cycles and cell debris was pelleted by centrifugation at 20,000 xg for 30 minutes at 4ºC. *E. coli* H16 was used for SAXS experiments.

### Protein purification

The cell-free reactions were diluted 5-10 fold with buffer A (50 mM Tris-HCl pH 7.5, 500 mM NaCl, 5 mM imidazole) before incubating it 1 h with 1.5 mL of Ni-resin (cOmplete™ His-Tag Purification Resin). The matrix was packed by gravity-flow and washed with buffer B (50 mM Tris-HCl pH 7.5, 1000 mM NaCl, 5 mM imidazole) and the target protein was eluted with buffer C (50 mM Tris-HCl pH 7.5, 150 mM NaCl, 250 mM imidazole). For SAXS measurements, this affinity chromatography step was carried out in an AKTA pure System (GE Healthcare) with a 5 mL Histrap® Excel column. Elution fractions were checked under UV light and fluorescent fractions were pooled, protease inhibitors were added (cOmplete EDTA-free protease inhibitor cocktail, Sigma Aldrich) and the sample was dialyzed against NMR buffer (20 mM BisTris-HCl pH 6.5, 150 mM NaCl) at 4°C using SpectraPor 4 MWCO 12-14 kDa dialysis tubing (Spectrum Labs). Dialyzed protein was then concentrated with 10 kDa MWCO Vivaspin centrifugal concentrators (3,500 xg, 4°C) (Sartorius). Protein concentrations were determined by means of fluorescence using an sfGFP calibration curve. Final NMR sample concentrations ranged from 4 to 15 µM. Protein integrity was analyzed by SDS-PAGE.

For both aggregation and SAXS experiments, an additional size-exclusion chromatography step, using a Superdex S200 10/300 column, was carried out. For aggregation measurements, this step was performed in aggregation buffer (50 mM sodium phosphate, pH 7.5, 150 mM NaCl) and for SAXS measurements in NMR buffer.

### NMR experiments and data analysis

All NMR samples contained final concentrations of 10% D_2_O and 0.5 mM 4,4-dimethyl-4-silapentane-1-sulfonic acid (DSS). ^15^N and ^13^C-HSQC experiments, in order to determine amide (^1^H_N_ and ^15^N) and aliphatic (^1^H_aliphatic_ and ^13^C_aliphatic_) chemical shifts, were performed at 293 K on a Bruker Avance III spectrometer equipped with a cryogenic triple resonance probe and Z gradient coil, operating at a ^1^H frequency of 700 MHz or 800 MHz. Spectra acquisition parameters were set up depending on the sample concentration and the magnet strength. All spectra were processed with TopSpin v3.5 (Bruker Biospin) and analyzed using CCPN-Analysis software^70^. Chemical shifts were referenced with respect to the H_2_O signal relative to DSS using the ^1^H/X frequency ratio of the zero point according to Markley *et al*.^71^.

Random coil chemical shifts were predicted using POTENCI, a pH, temperature and neighbor corrected IDP library (http://nmr.chem.rug.nl/potenci/)^37^. Secondary chemical shifts (SCS) were obtained by subtracting the predicted value from the experimental one (SCS=δ_exp_-δ_pred_). For better reliability of the results regarding possible referencing errors, we used the combined C_α_ and C_β_ secondary chemical shifts (SCS(C_α_)-SCS(C_β_)).

### ^19^F-NMR experiments

All NMR samples were first concentrated up to a ca. 200 µl volume using Vivaspin centrifugal concentrators (Sartorius) with a 5 kDa cutoff at 4°C. 0.1 µl of a trimethylsilylpropanoic acid (TMSP) solution for chemical shift referencing and 10 µl of D_2_O were added before NMR measurement. All ^19^F NMR experiments were performed on a Bruker Avance III HD spectrometer operating at a ^1^H and ^19^F frequencies of 600.13 MHz and 564.69 MHz, respectively, equipped with a CP-QCI-F cryoprobe with ^19^F cryo-detection. All ^19^F 1D experiments were performed at 293.0 K with ^1^H decoupling during acquisition using waltz16 composite pulse decoupling. An acquisition time of 0.58 s, spectral width of 100.6 s and relaxation delay of 1.0 s was used for all samples, except for the sample fluorinated at Q20, where an acquisition time of 0.29 s and a relaxation delay of 0.5 s was used. Concatenated 1D ^19^F spectra of 128 transients each were acquired in order to monitor any spectral changes over time. Signal averaging was then performed up to a time point before significant spectral changes over time could be detected. Final number of transients varied were 77824, 49152, 39680 and 53760 for samples fluorinated at Q20, Q21, Q40 and Q60, respectively. ^19^F spectra were referenced to the ^1^H signal of TMSP using the unified chemical shift scale.

### Model building and chemical shift ensemble optimization

Ensemble models for the two families capturing the conformational influences of the flanking regions, N→C and N C, were constructed with the algorithm described in reference^38^, which uses a curated database of three-residue fragments extracted from high-resolution protein structures. The model building strategy consecutively appends residues, which are considered to be either fully disordered or partially structured. For fully disordered residues, amino acid specific ϕ/ψ angles defining the residue conformation are randomly selected from the database, disregarding their sequence context. For partially structured residues, the nature and the conformation of the flanking residues are taken into account when selecting the conformation of the incorporated residue. Steric clashes are tested at each step, and a backtracking strategy is applied to solve possible conflicts (see detailed explanation of the algorithm in reference^38^).

Two families of ensembles were built. For the first family (N→C ensembles), starting with the ^10^AFESLKS^16^ region of N17 as partially structured, multiple ensembles of 5,000 conformations were built by successively including an increasing number of glutamines in the poly-Q tract (from F17 to Q63) as partially structured, while the rest of the chain was considered to be fully disordered. An equivalent strategy was followed for the second family of ensembles (N C ensembles) for which glutamines were considered successively as partially structured from the poly-P tract (from Q63 to Q18). Note that in the partially structured building strategy secondary structural elements are propagated due to the conformational influence of neighboring residues. Two tripeptide databases were used to generate the conformational ensemble models. Both were constructed from the protein domains in the SCOP (Structural Classification of Proteins)^72,73^ repository filtered to 95% sequence identity. An “unfiltered” tripeptide database was built disregarding secondary structure content, and “coil” database included tripeptides not participating in α-helices or β-strands. For the N→C ensembles, the best results were obtained when using the “unfiltered” and “coil” databases to sample the partially-structured and the fully disordered sections, respectively. For the N←C ensembles, the “coil” database yielded the best results. For the resulting 47 ensembles of each family, and after building the side chains with the program SCWRL4^39^, averaged Cα chemical shifts were computed with SPARTA+^40^, and used for the optimization. The optimized ensemble model of H46 was built by reweighting the populations of the pre-computed ensembles, minimizing the error with respect to the experimental Cα CSs. In order to capture the influence of the flanking regions, glutamines within the tract were divided into two groups: those influenced by N17 and those influenced by the poly-P tract, whose chemical shifts were fitted with the N→C and N←C ensembles, respectively. The limit between both families was systematically explored by computing the agreement between the experimental and optimized CSs through a χ^2^ value. An optimal description of the complete CS profile was obtained when Q55 was chosen as the last residue structurally connected with N17. Finally, an ensemble of 11,000 conformations was built using the optimized weights and it was used to derive secondary structure population using SS-map^43^ and to analyze the SAXS data.

### SAXS data measurement and analysis

The SAXS data for H16 were collected at the SWING beamline at the SOLEIL synchrotron, France, equipped with an Eiger 4M detector with a sample-to-detector distance of 1.5 m^74^. The data for H46 were collected at EMBL-bioSAXS-P12 beamline at PETRAIII, Hamburg, Germany equipped with a Pilatus 6M detector with a sample-to-detector distance of 3 m^75^. The parameters used for SAXS data collection are given in Table S2. All the data were collected in SEC-SAXS mode with an in-line Superdex 200 Increase 10/300 GL column (GE Healthcare). Both proteins were concentrated to 8 mg/mL and centrifuged at 20,000 x g immediately before injecting the protein onto the column. 80 µL of the sample were injected into the column and the flow rate was maintained at 0.5 mL/min. The initial data processing steps including masking and azimuthal averaging were performed using the program FOXTROT^76^ for H16 and SASFLOW pipeline^75^ for H46. The resulting 1D profiles were analyzed using CHROMIXS^77^ from ATSAS suite to select the frames corresponding to sample and buffer and perform buffer subtraction. The final buffer subtracted and averaged SAXS profiles were analyzed using ATSAS 2.8 software package^78^, including AUTORG for calculating the radius of gyration and calculation of extrapolated value of radius of gyration (*R*_*g*_), GNOM^79^ for calculation of pairwise distance distribution profiles and DATBAYES^80^ for calculation of molecular weight by Bayesian estimate from four approaches. The ensemble optimization approach (EOM) was used to select sub-ensembles that collectively describe the SAXS data. The program RanCH was first used to join each of the conformations of H16 and H46 (generated as described above) to the modelled structure of sfGFP and the hexahistidine tag used for purification. GAJOE was then used to find a sub-ensemble from this pool which collectively describes the SAXS data^41,42^. The graphical representations were generated using the program VMD^81^.

### Molecular dynamics simulations

We performed Gaussian Accelerated Molecular Dynamics (GaMD)^48^ simulations to explore the conformational landscape and the secondary structure propensities of a fragment of httex1 consisting of N17, 46 glutamines and 5 prolines. We used ff03ws-STQ^50^ force field, which is adapted to proteins with low-complexity sequences (obtained from https://bitbucket.org/jeetain/all-atom_ff_refinements/src/master/). We chose an extended conformation built using the protocol described earlier in the model building section as the starting structure and prepared the simulation system using tools available with GROMACS 2020.5. This included addition of hydrogens, solvation and addition of ions (Na^+^ and Cl^-^) to neutralize the system and set the final salt concentration to 0.15 M. Thereafter, the system was converted to AMBER format using ParmEd tool in AMBER20. At this stage, hydrogen mass repartitioning was also done to allow a time step of 4 fs. We used periodic boundary conditions and restrained the bonds containing hydrogen atoms using SHAKE algorithm. Particle Mesh Ewald summation (PME) was used to calculate electrostatic interaction with a cut-off of 9 Å on the long-range interactions. The system was energy minimized for 5000 steps. This was followed by an NVT (constant number, volume and temperature) equilibration for 5 ns and further equilibration in NPT (constant number, pressure and temperature) for 10ns. For all the simulations, the temperature was maintained at 293 K and for NPT simulations the pressure was maintained at 1 atm. This was followed by a GaMD equilibration stage which consisted of a classical MD simulation of 40 ns during which the potential statistics to calculate GaMD parameters were collected followed by a 120 ns long equilibration during which the boost was added and updated. Finally, 8 independent simulations with an aggregate simulation time of ∼ 20 µs were launched using the boost parameters obtained in the equilibration stage. The simulations were run in “dual-boost” mode and the reference energy was set to the lower bound (E=V_max_). The average and standard deviation of the potential energy were calculated every 2ns (number of steps ≈ 4*system size).

### Aggregation experiments

Time-dependent aggregation of H46 variants was followed with SDS-PAGE analysis as previously described, with minor modifications^52^. 15 µM wild-type-, LKGG- and LLLF-H46 samples, prepared in aggregation buffer (50 mM sodium phosphate, pH 7.5, 150 mM NaCl), were incubated at 37°C for 48 h, without shaking. 10 µL-aliquots were extracted at different time intervals, immediately mixed with denaturing buffer (125 mM Tris-HCl, pH 6.8, 20% glycerol, 4% SDS, 200 mM dithiothreitol, 0.05% bromophenol blue), incubated for 10 min at 95°C, and frozen at −20 °C until analysis on Bolt™ 4–12% Bis-Tris Plus gels (Invitrogen). The gels were washed in water, stained with Instant Blue Coomassie Protein Stain (Abcam), and visualized using a Gel Doc™ Ez Imager (Bio-Rad). The amount of SDS-soluble species trapped in the stacking gel was quantified using the Image Lab 5.1 software. The percentages of soluble protein were referenced to time 0 values, and plotted against time. The plots were fitted using GraphPad Prism Software. For each protein variant, at least two independent experiments, with three replicates at each time point, were recorded.

### Atomic force microscopy (AFM) and total internal reflection fluorescence (TIRF)

Correlative AFM-TIRF microscopy was developed in-house^82^. AFM images were acquired using a Nanowizard 4 (JPK Instruments, Bruker) mounted on a Zeiss inverted optical microscope and equipped with a Vortis-SPM control unit. A custom-made TIRF microscope was coupled to the AFM using a LX 488-50 OBIS laser source (Coherent). We used an oil immersion objective with a 1.4 numerical aperture (Plan-Apochromat 100x, Zeiss). Fluorescence was collected with an EmCCD iXon Ultra897 (Andor) camera. The setup includes a 1.5x telescope to obtain a final imaging magnification of 150-fold, corresponding to a camera pixel size of 81.3 nm. An ET800sp short pass filter (Chroma) was used in the emission optical path to filter out the light source of the AFM optical beam deflection system. The excitation laser wavelength was centered at 488 nm and the power was measured before the objective with a PM100 energy meter (purchased from Thorlabs) and was optimized in all the experiments in the range of 1-5 *µ*W. Fluorescence images were acquired using an ET525/50 nm (Chroma) emission filter and an acousto-optic tuneable filter (AOTFnc-400.650-TN, AA opto-electronics) to modulate the laser intensity. Fluorescence images were obtained by averaging 150 individual images, each acquired over 50 ms as exposure time.

AFM images were collected in liquid environment (Dulbecco’s PBS named D-PBS) using the quantitative-imaging (QI) mode. Each image was acquired with 256×256 lines/pixels and the following scan size: 5 µm × 5 µm, 2.5 µm × 2.5 µm and 1 µm × 1 µm. Typical force *versus* distance curves were recorded with a tip approach speed ranging from 10 µm/s to 30 µm/s and an oscillation amplitude (Z length) of 100 nm or 150 nm, adjusted depending on the height of the aggregates. The maximal force exerted in each pixel was set to 100-150 pN and optimized during the image acquisition. We used MSNL-D and MSNL-E (Bruker) AFM probes with resonances in liquid of ≈ 2 kHz and 10 kHz, respectively, and nominal spring constants of 0.03 N/m and 0.1 N/m. MSNL cantilevers have a sharp tip radius (≈ 2 nm), which is ideal for high-resolution imaging. The inverse optical lever sensitivity was calibrated with the acquisition of a force *versus* distance curve on the glass coverslip whereas the cantilever stiffness was calibrated using thermal method^83^.

Samples for correlative AFM and TIRF were prepared on circular glass coverslips (2.5 cm, 165 µm thick, purchased from Marienfeld). Coverslips were cleaned with a 15 min cycle of sonication with ultrasounds in 1M KOH, rinsed 20 times with deionized water and finally with a second cycle of sonication in deionized water. Fibrils were then deposited on the clean glass coverslips and let dry before being immersed in D-PBS for AFM imaging.

## Acknowledgements

The authors thank Gottfried Otting for providing the BL21 (DE3) Star::RF1-CBD3 strain. This work was supported by the European Research Council under the European Union’s H2020 Framework Programme (2014-2020) / ERC Grant agreement n° [648030] awarded to PB, ANR-17-CE11-0022-01 awarded to NS, and Labex EpiGenMed, an « Investissements d’avenir » program (ANR-10-LABX-12-01). The CBS is a member of France-BioImaging (FBI) and the French Infrastructure for Integrated Structural Biology (FRISBI), 2 national infrastructures supported by the French National Research Agency (ANR-10-INBS-04-01 and ANR-10-INBS-05, respectively). AU is supported by a grant from the Fondation pour la Recherche Médicale (SPF20150934061). DS acknowledge a grant from the Métropole Européenne de Lille (PUSHUP). Géraldine Levy, Université de Lille, is thanked for help with sample preparation and the ^19^F-NMR experiments. This work benefited from the HPC resources of the CALMIP supercomputing center under the allocations 2016-P16032 and 2021-P21043. The 600 MHz spectrometer for ^19^F NMR measurements is funded by the Nord Region Council, CNRS, Institut Pasteur de Lille, the European Community (ERDF), the French Ministry of Research and the Université de Lille and by the CTRL CPER cofunded by the European Union with the European Regional Development Fund (ERDF), by the Hauts-de-France Regional Council (contract n°17003781), Métropole Européenne de Lille (contract n°2016_ESR_05), and French State (contract n°2017-R3-CTRL-Phase1). The authors thank the SWING beamline at SOLEIL synchrotron, Saint-Aubin, France (proposal 20181386) and P12 beamline at PETRAIII, Hamburg, Germany for beamtime allocation to the project and assistance during data collection.

## Author Contributions

P.B. conceived the project. C.A.E.R, A.U., A.S., A.B., D.S., N.S. and P.B. designed experiments. C.A.E.R, A.U., A.S., M.P., A.M., A.E., A.F., X.L.L., Z.D.S., L.C., A.T., F.A. and D.S. performed experiments. R.E.S., P.E.M., A.B., J.C., N.S. and P.B. supervised experiments. C.A.E.R., A.U., A.S. and P.B. wrote the manuscript with the help of all the co-authors.

## Data Availability Statement

The datasets generated during and/or analysed during the current study are available from the corresponding author on reasonable request.

