## Supplemental information for "The structure of pathogenic huntingtin exon-1 defines the bases of its aggregation propensity"

**Table S1.** Glutamine chemical shifts measured for H46 and sample concentrations.

| Residue | N | HN | Cα | Cβ1 / Cβ2 | Cγ | Nε | Hα | Hβ1 / Hβ2 | Hγ1 / Hγ2 | Hε1 / Hε2 | Conc. [μM] |
|---|---|---|---|---|---|---|---|---|---|---|---|
| Q18 | 120.45 | 8.2978 | 57.279 | 29.774 / 29.762 | 33.997 | 111.92 | 4.125 | 2.116 / 2.051 | 2.384 | 6.862 / 7.504 | 18.8 |
| Q20 | 120.628 | 8.282 | 57.276 | 29.003 / 28.977 | 33.992 / 33.98 | 112.15 | 4.183 | 2.055 / 2.111 | 2.419 / 2.398 | 6.869 / 7.462 | 9 |
| Q21 | 120.338 | 8.285 | 57.27 | 28.965 / 29.002 | 33.919 / 33.927 | 112.78 | 4.126 | 2.058 / 1.964 | 2.433 / 2.393 | 6.86 / 7.394 | 17.1 |
| Q24 | 120.51 | 8.312 | 57.302 | 29.013 / 28.962 | 33.926 / 33.9 | 112.11 | 4.214 | 2.136 / 2.394 | 2.438 / 2.394 | 6.865 / 7.525 | 8 |
| Q28 | 120.731 | 8.339 | 57.048 | 29.044 / 29.015 | 33.914 / 33.919 | 112.06 | 4.239 | 2.133 / 2.071 | 2.428 / 2.396 | 6.853 / 7.509 | 4 |
| Q32 | 120.756 | 8.364 | 57.177 | 29.019 / 29.006 | 33.89 / 33.892 | 112.192 | 4.231 | 2.082 / 2.141 | 2.435 / 2.339 | 6.88 / 7.539 | 5 |
| Q36 | 120.806 | 8.371 | 57.024 | 29.037 / 28.957 | 33.909 / 33.925 | 112.22 | 4.235 | 2.133 / 2.079 | 2.437 / 2.397 | 6.88 / 7.54 | 5 |
| Q40 | 120.842 | 8.376 | 57.015 | 28.962 / 29.061 | 33.908 / 33.851 | 112.24 | 4.236 | 2.135 / 2.072 | 2.437 / 2.399 | 6.88 / 7.541 | 3.5 |
| Q44 | 120.908 | 8.381 | 57.009 | 29.084 / 29.023 | 33.952 / 33.924 | 112.25 | 4.24 | 2.066 / 2.141 | 2.395 / 2.431 | 6.882 / 7.541 | 6 |
| Q48 | 120.929 | 8.389 | 56.917 | 29.116 / 29.079 | 33.899 / 33.916 | 112.19 | 4.251 | 2.128 / 2.073 | 2.411 / 2.399 | 6.882 / 7.542 | 4.5 |
| Q52 | 121.056 | 8.404 | 56.721 | 29.183 / 29.131 | 33.896 | 112.33 | 4.258 | 2.05 / 2.117 | 2.402 | 6.879 / 7.542 | 5 |
| Q56 | 121.114 | 8.411 | 56.415 | 29.241 / 29.203 | 33.865 | 112.38 | 4.274 | 2.031 / 2.119 | 2.397 | 6.884 / 7.544 | 2.6 |
| Q60 | 121.436 | 8.438 | 56.07 | 29.423 / 29.267 | 33.883 | 112.5 | 4.303 | 2.002 / 2.111 | 2.39 | 6.892 / 7.557 | 5 |
| Q61 | 121.562 | 8.449 | 55.966 | 29.39 / 29.393 | 33.833 | 112.525 | 4.313 | 2.097 / 1.996 | 2.377 | 6.892 / 7.56 | 4 |
| Q62 | 121.934 | 8.47 | 55.765 | 29.583 / 29.526 | 33.804 | 112.7 | 4.315 | 1.977 / 2.085 | 2.366 | 6.887 / 7.567 | 8 |
| Q63 | 123.131 | 8.458 | 53.604 | 28.951 / 28.844 | 33.466 | 112.64 | 4.607 | 2.086 / 1.932 | 2.403 | 6.879 / 7.543 | 5 |
| Q75 | 120.587 | 8.434 | 55.39 | 29.721 / 29.668 | 33.855 | 113.042 | 4.317 | 2.051 / 1.949 | 2.37 | 6.91 / 7.612 | 5.2 |
| Q91 | 122.041 | 8.516 | 53.498 | 29.007 / 28.989 | 33.492 | 113.134 | 4.592 | 2.082 / 1.924 | 2.427 | 6.92 / 7.597 | 5 |

**Table S2.** SAXS table.

| **Data-collection parameters** | **H16** | **H46** |
|---|---|---|
| Instrument | SWING (SOLEIL) | P12 (EMBL PETRA III) |
| Beam geometry | Point collimation | Point collimation |
| Detector | EigerX4M | Pilatus 6M |
| Wavelength (Å) | 1.0332 | 1.2398 |
| Sample to detector distance | 1.5 m | 3.0 m |
| Measured q range ($Å^{-1}$) | 0.004-0.56 | 0.002-0.65 |
| Temperature (K) | 283 | 283 |
| Exposure time (ms) | 990 ms *2100 frames | 245 ms * 1680 frames |
| Concentration (mg $mL^{-1}$) | 8 | 8 |
| Calculated monomeric ($M_r$) from sequence | 39.1 | 42.9 |
| **Software employed** | | |
| Primary data reduction | FOXTROT | SASFLOW |
| Data processing | CHROMIXS | CHROMIXS |
| Guinier Analysis | PRIMUS QT | PRIMUS QT |
| IFT Analysis | GNOM 4.6 | GNOM 4.6 |
| Ensemble Fitting | EOM | EOM |
| Three-dimensional graphics representations | PyMOL, VMD | PyMOL, VMD |
| **Structural parameters†** | | |
| *I(0)* (AU) (from *P(r)*) | 0.02 | 0.02 |
| $R_g$ (Å) (from *P(r)*) | 33.5 | 43.5 |
| *I(0)* (AU) (from Guinier) | 0.02 ± 9e-05 | 0.02 ± 1.7e-04 |
| $R_g$ (Å) (from Guinier) | 32.9 ± 0.2 | 41.5 ± 0.5 |
| $D_{max}$ (Å) | 126 | 148 |
| **Molecular-mass determination** | | |
| Molecular mass ($M_r$) [Bayesian Estimate] | 39.3 | 58.1 |
| **SASBDB Entry** | - | - |

### FIGURE S1

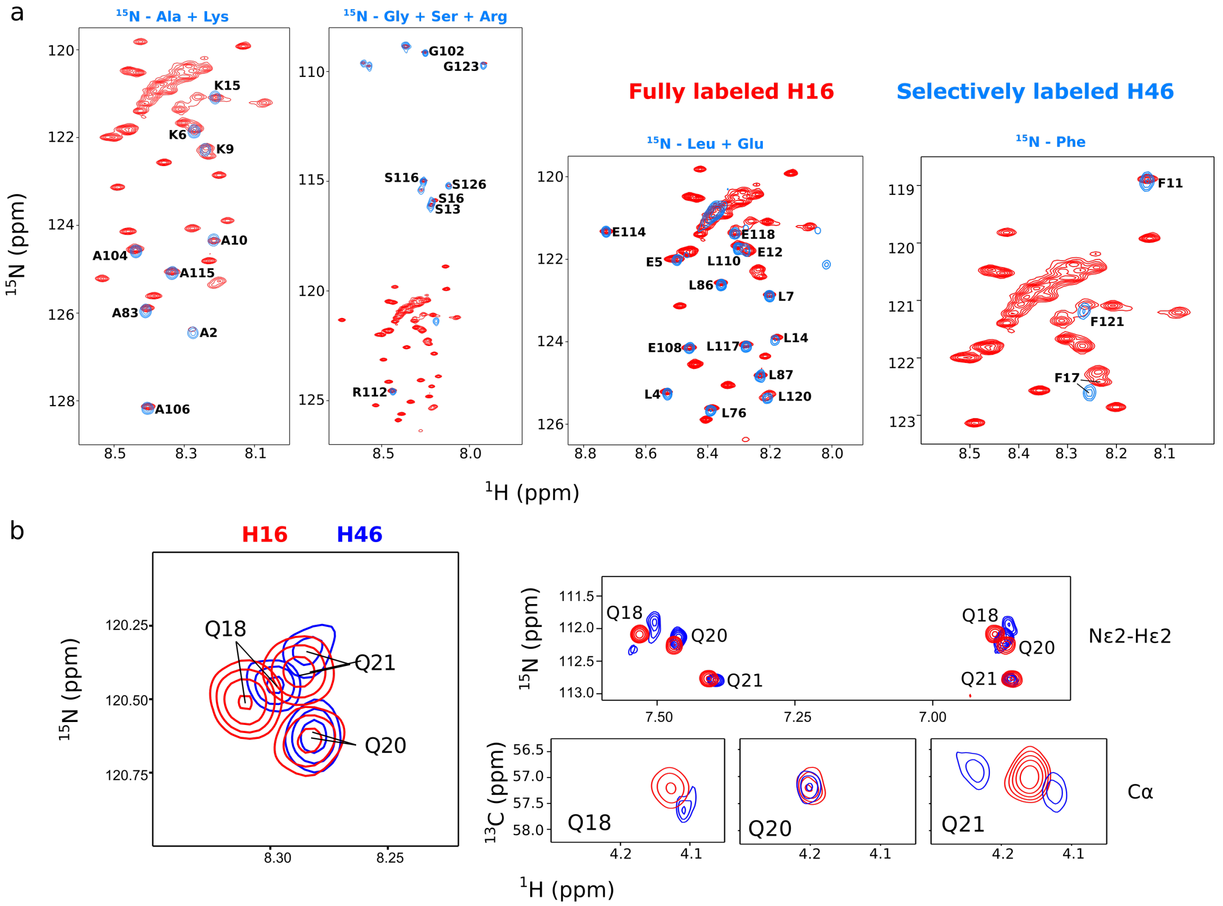


**Figure S1. Comparison of H46 and H16 by NMR. (a)** Overlay of the $^{15}$N-HSQC spectra of fully labeled H16 (red) with selectively labeled samples of H46 (blue). **(b)** Zoom of the $^{15}$N-HSQC and $^{13}$C-HSQC overlay of H16 and H46 SSIL spectra showing the poly-Q NH, NHε and Cα regions of Q18, Q20 and Q21.

**FIGURE S2**

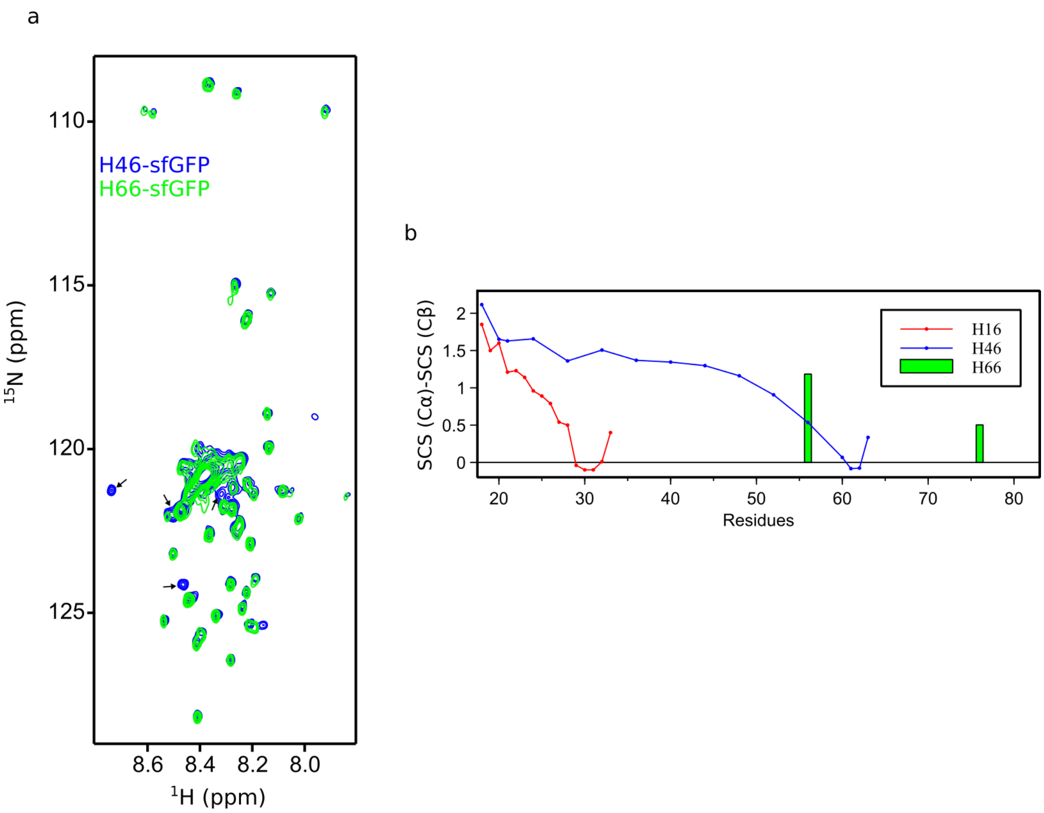


**Figure S2. Comparison of H66 with H16 and H46 by NMR. (a)** Overlay of the $^{15}$N-HSQC spectra of fully labeled H66 (green) with fully labeled H46 (blue). Black arrows indicate peaks corresponding to glutamates, which were not labeled in the H66 sample. **(b)** Comparison of the SCS profiles of H16 (red), H46 (blue) and the values measured for Q56 and Q76 in H66 (green).

FIGURE S3

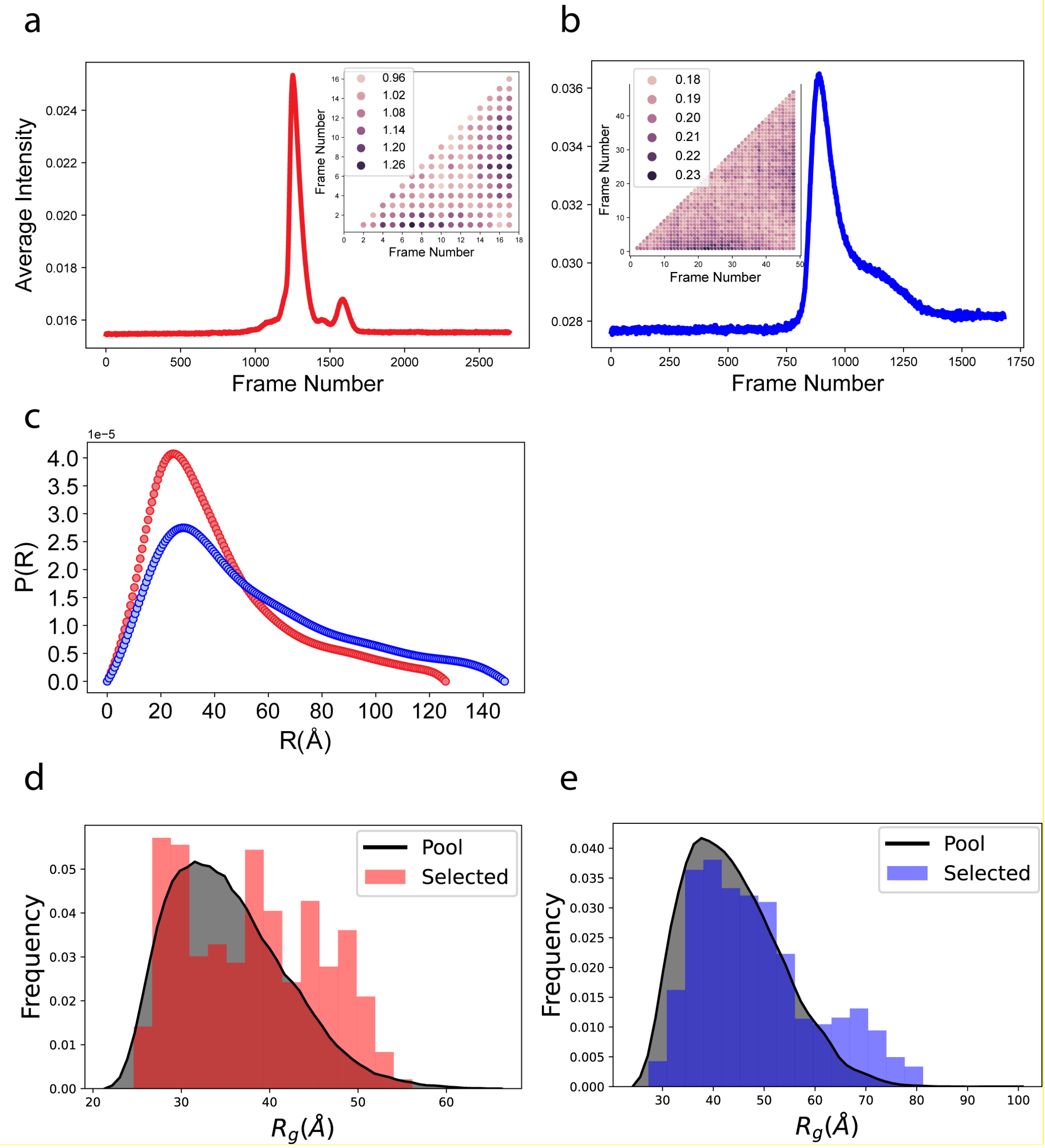


**Figure S3. SAXS analyses of H46 and H16.** The plots of average intensity *vs.* frame number obtained from SEC-SAXS for H16 **(a)** and H46 **(b)**. The insets show all-*vs*-all $\chi^2$ comparison for the frames selected for further processing. The small values of $\chi^2$ show that the selected frames were very similar to each other. **(c)** Pairwise distance distribution functions, *P(r)*, obtained for H16 (red) and H46 (blue) by indirect Fourier transformation of the SAXS data. **(d)** and **(e)** show the $R_g$ distribution of the pool (gray, filled) and the selected sub-ensemble by EOM for H16 (red) and H46 (blue), respectively.

#### FIGURE S4

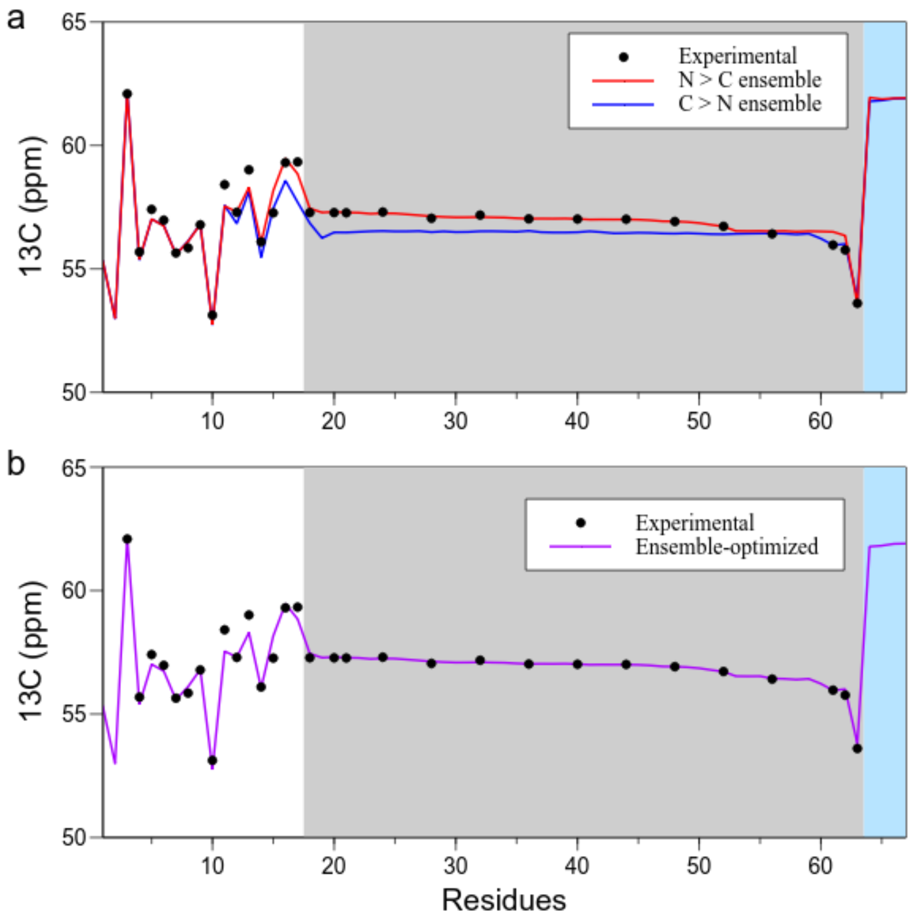


**Figure S4. Chemical shift based ensemble refinement of H46. (a)** Experimental (black) *vs.* ensemble-optimized (red for N→C and blue for N←C ensembles) chemical shifts for H46. **(b)** The final optimized ensemble was built using the M1-Q55 and the Q55-P113 fragments of the N→C and N←C ensembles, respectively. The poly-Q tract is shaded in gray. Notice that we have measured the chemical shiftd for 16 glutamines of the 46-glutamine long tract. No experimental data for prolines are available. The PRR is shaded in blue.

#### FIGURE S5

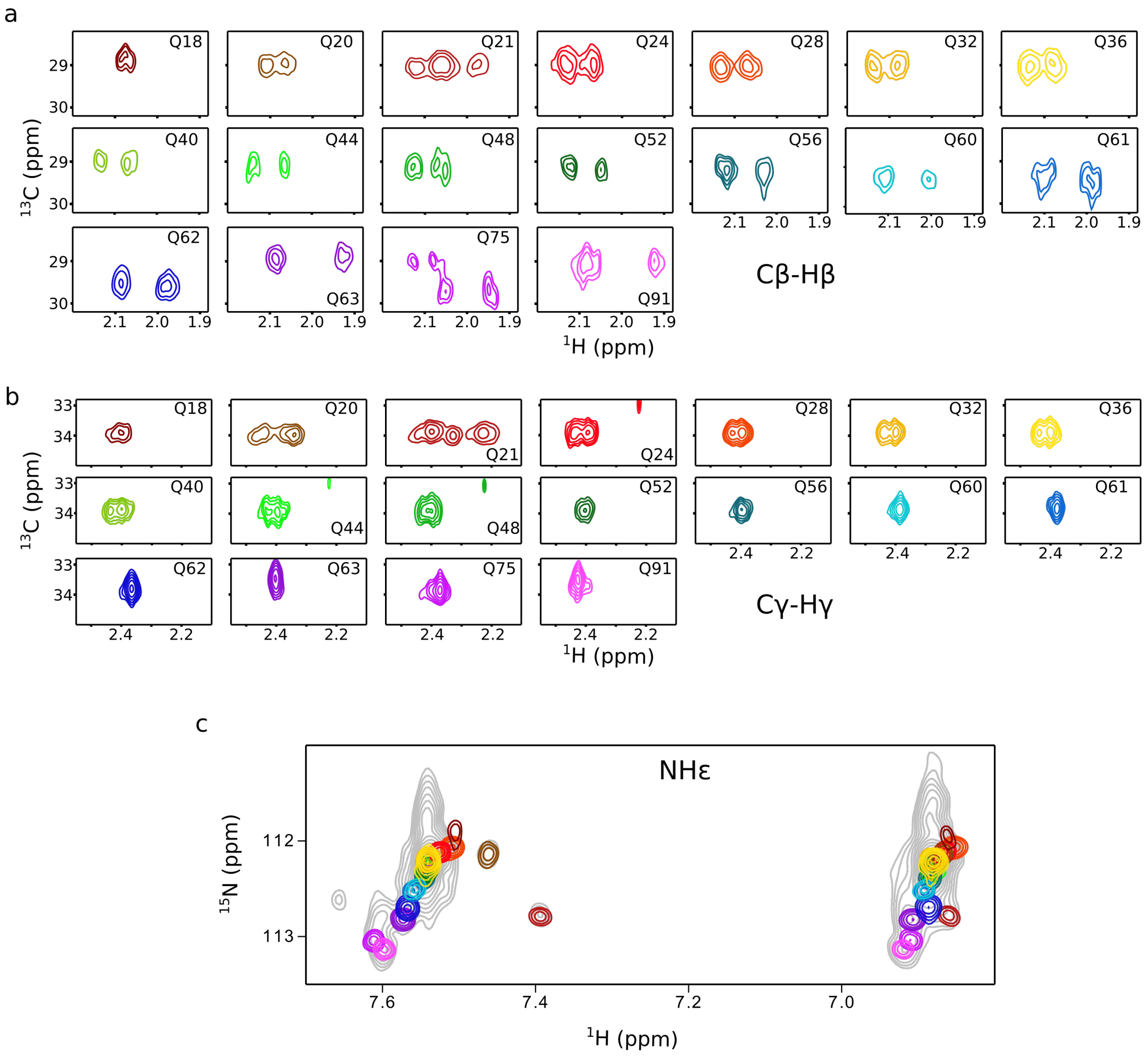


**Figure S5. Side chain NMR scanning. (a)** Cβ-Hβ and **(b)** Cγ-Hγ zooms of the $^{13}$C-HSQC spectra for all glutamines scanned in H46. **(c)** Zoom of the $^{15}$N-HSQC spectra showing the side chain $N\varepsilon H_2$ region. The color code is equivalent to the one used in Figures 1 and 3 in the main text.

**FIGURE S6.**

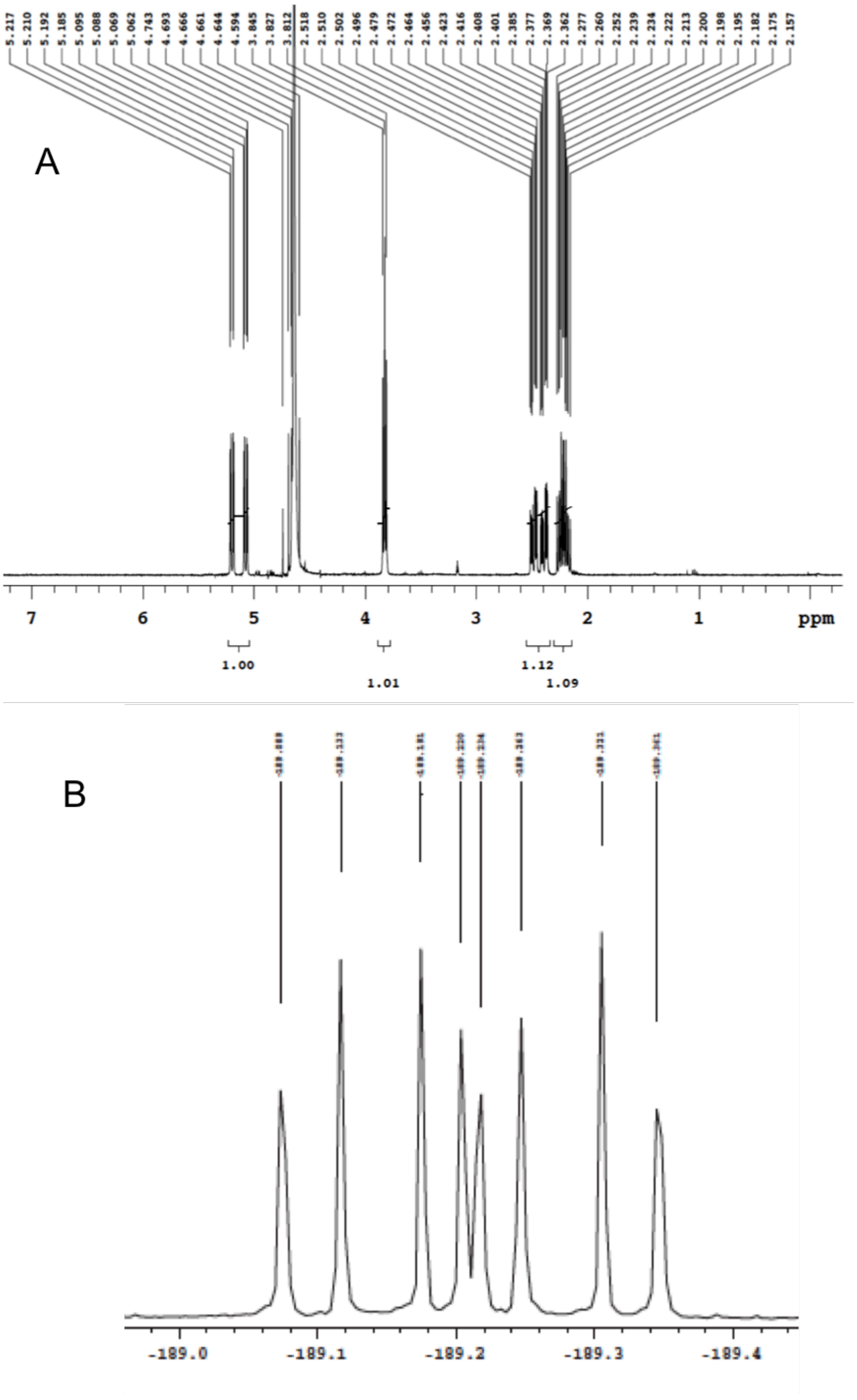


**Figure S6.** $^1$H- (**A**) and $^{19}$F-NMR (**B**) spectra of 2S,4R-fluoroglutamine (4F-Gln).

#### FIGURE S7

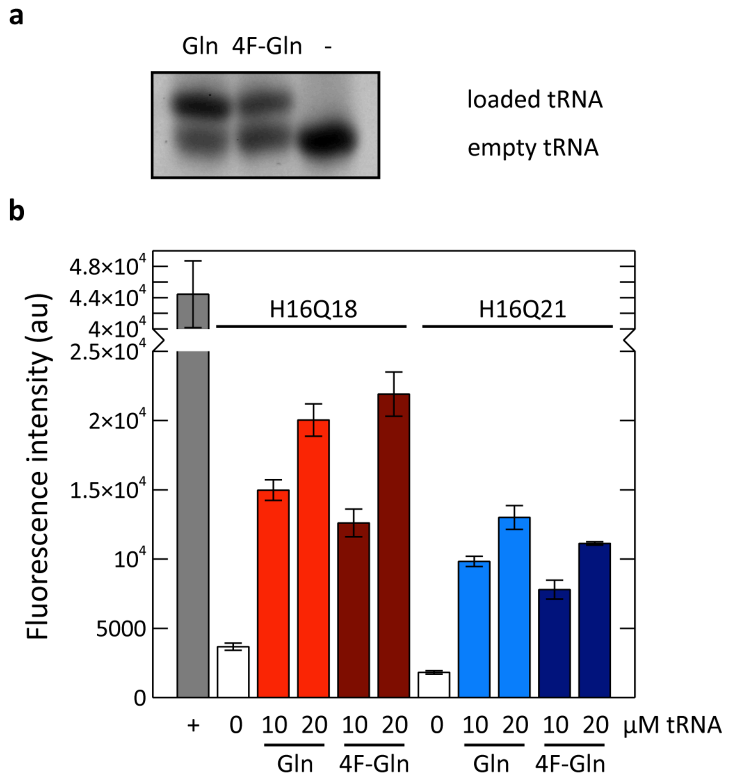


**Figure S7. Labeling of httex1 with fluorinated glutamines. (a)** Enzymatic loading of suppressor $tRNA_{CUA}$ with canonical (Gln) or 2S,4R-fluoroglutamine (4F-Gln). Upper and lower bands correspond to loaded and unloaded suppressor $tRNA_{CUA}$, respectively. A negative control of empty $tRNA_{CUA}$ is shown in the third lane. **(b)** Endpoint fluorescence for cell-free suppression reactions of H16 with a stop codon at position Q18 or Q21 when titrating with increasing concentrations of both Gln-$tRNA_{CUA}$ and 4F-Gln-$tRNA_{CUA}$. "+" indicates a positive control, a cell-free reaction of H16 without any amber stop codon.

**FIGURE S8**

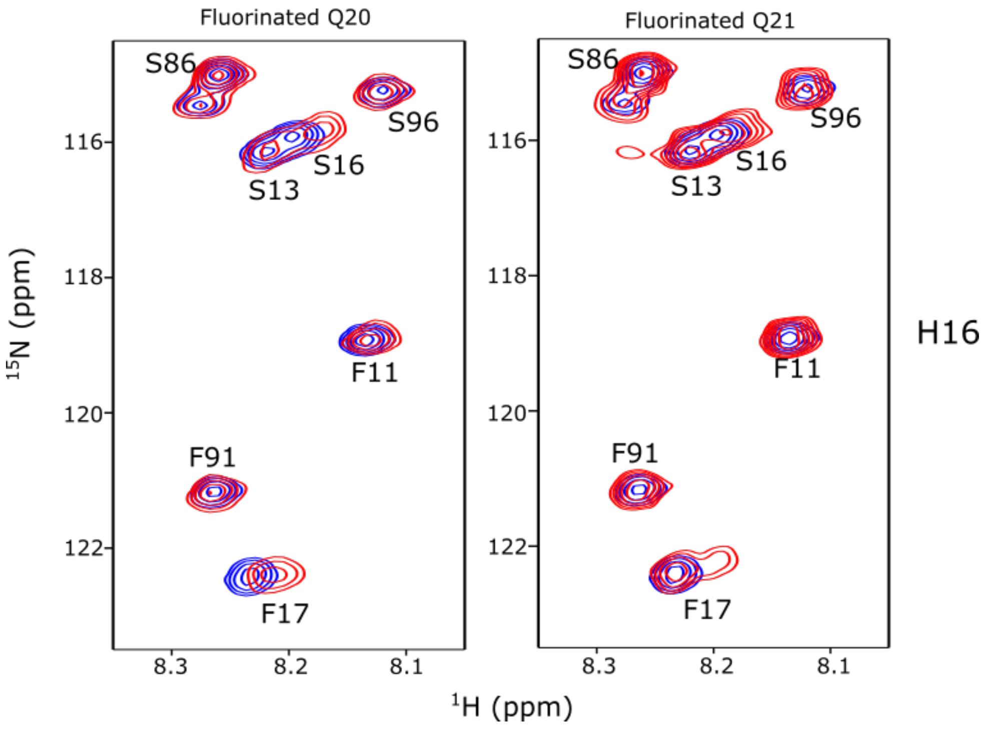


**Figure S8. NMR data of fluorinated H16.** $^{1}$H-$^{15}$N-HSQC spectra of H16 samples labeled with both $^{15}$N-Ser and/or $^{15}$N-Phe. In red, spectra of samples with fluorinated glutamine at position Q20 (right panels) or Q21 (left panels). Non-fluorinated samples are colored in blue.

**FIGURE S9.**

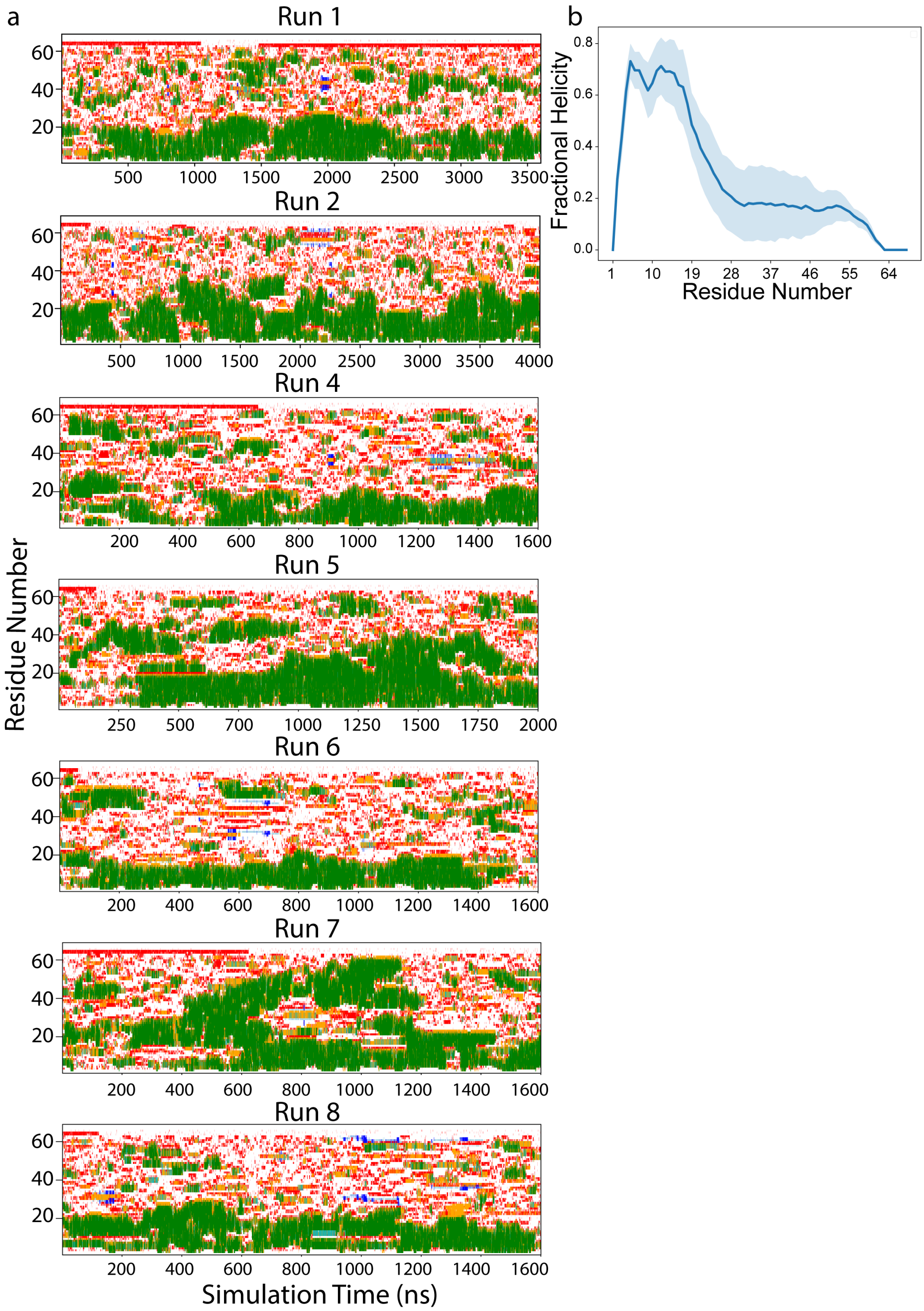


**Figure S9. MD simulations of H46. (a)** The per-residue secondary structure plot for the seven of the eight independent trajectories. The same plot for 'Run 3' is shown in Fig. 5a. **(b)** The weighted average of reweighted fractional helicity calculated after concatenating the 8 trajectories with an aggregated time of ≈20 μs. The shaded region shows the weighted standard deviation based on the number of frames for each trajectory.

**FIGURE S10.**

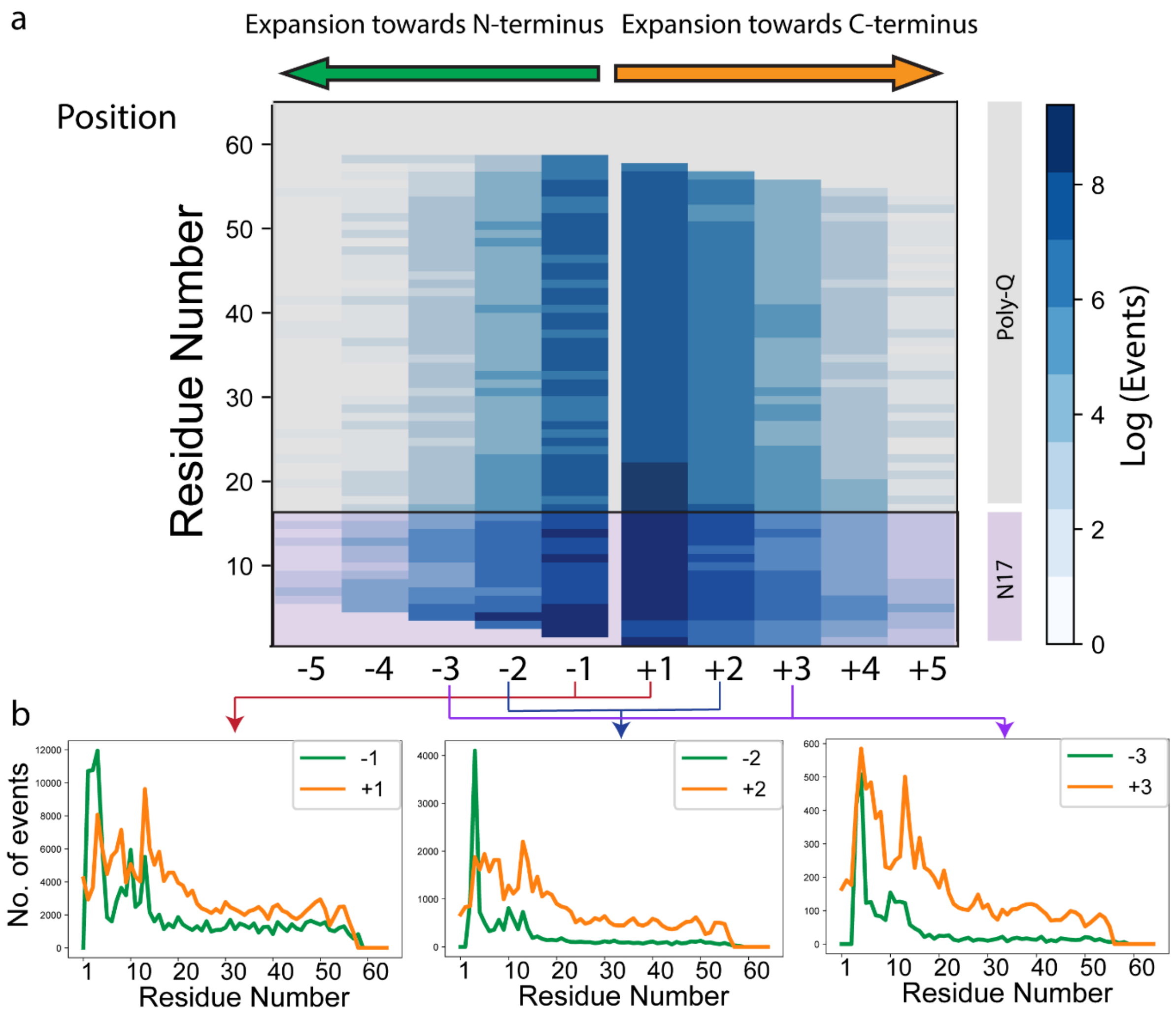


**Figure S10**. **Helicity propagation in H46 MD trajectories. (a)** Plot showing the number of times (in log scale) an α-helix expands 1 to 5 residues towards C-terminus (+1 to +5) or N-terminus (-1 to -5) from each of the residues of N17 and poly-Q. For all the positions (1 to 5), the number of events is greater in direction of C-terminus than the N-terminus indicating a higher propensity of helix propagation from N- to C-terminus. The comparison of number of events for expansion of helix by 1, 2 and 3 residues towards N or C-terminus is presented in **(b)** for further clarity. These plots show that in httex1 helical segments are more prone to expand towards the C-terminus than towards the N-terminus. Additionally, the preference for directional growing increases with the extent of propagation, *i.e.*, the relative difference in the number of events where a helix is propagated 3 residues towards C-terminus and N-terminus is larger than propagation for 2 residues, which is in turn larger than propagation by 1 residue.

**FIGURE S11**

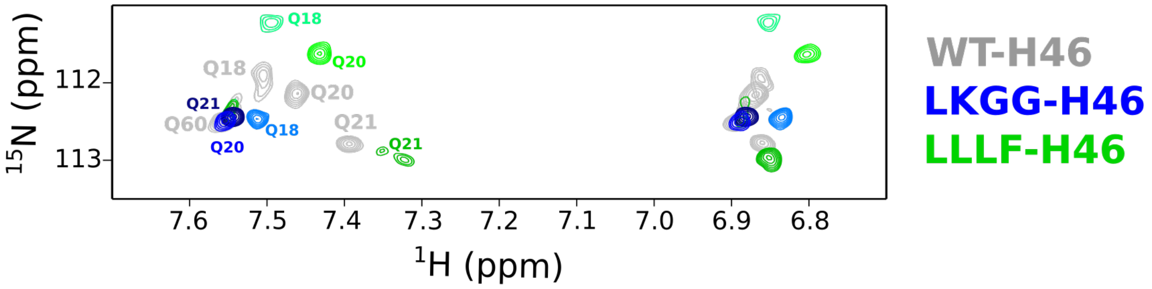


**Figure S11:** Zoom of the NεH$_2$ regions for the Q18, Q20 and Q21 SSIL samples for H46 (gray), LKGG-H46 (blue) and LLLF-H46 (green). The H46 Q60 signals are shown as an example of a glutamine adopting a random-coil conformation.

**FIGURE S12.**

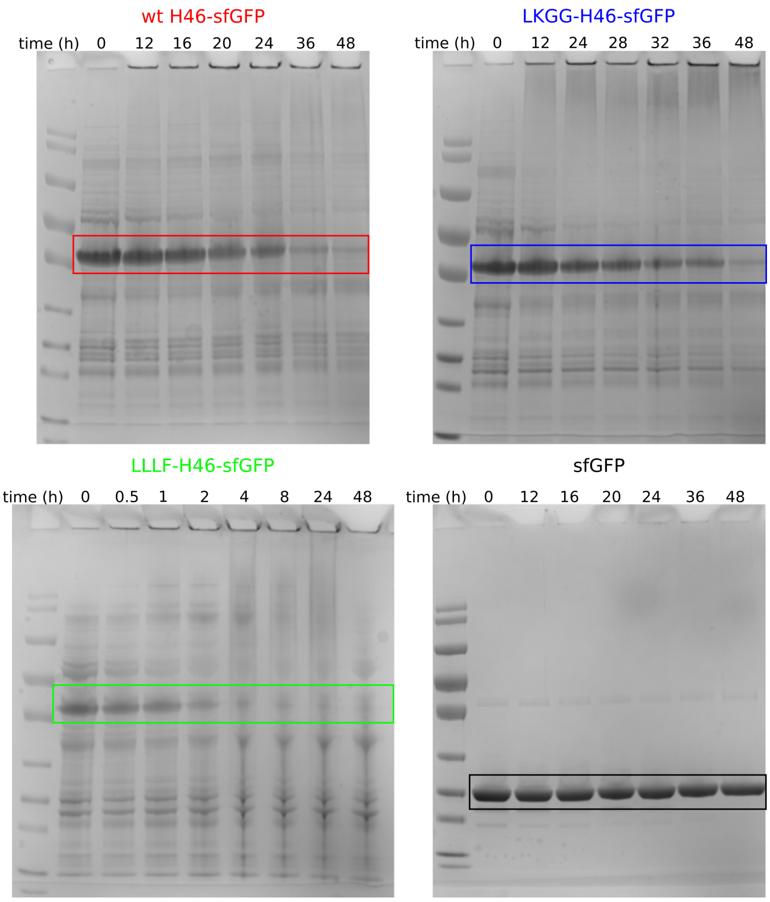


**Figure S12. Aggregation propensity of H46 variants.** SDS-PAGE for H46, LKGG-H46, LLLF-H46 and sfGFP along one of the aggregation experiments.

**FIGURE S13.**

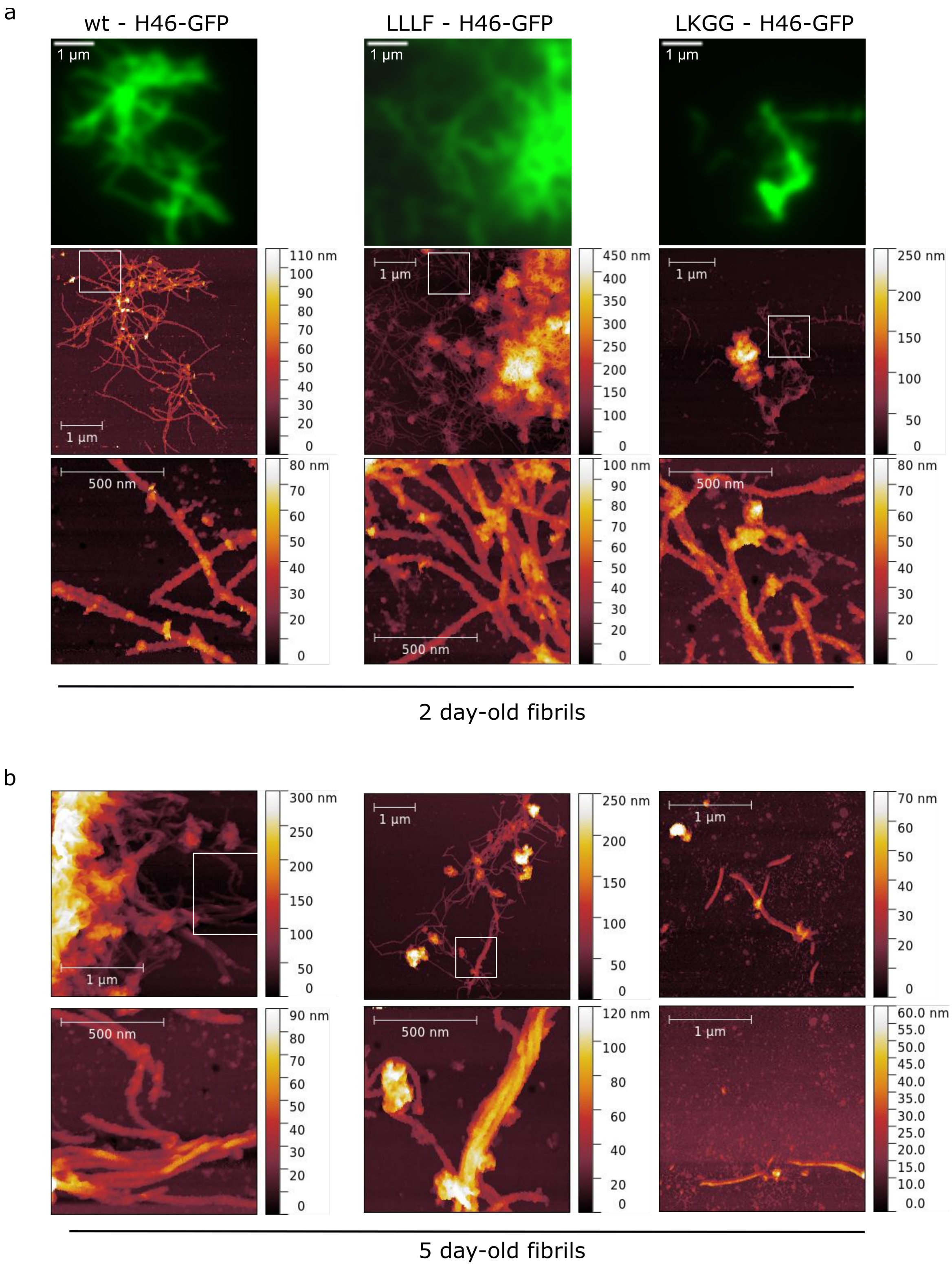


**Figure S13. (a)** Fluorescence microscopy (upper panels) and AFM (middle and lower panels) images of 2-day-old fibrils of H46, LLLF-H46 and LKGG-H46. Each fluorescence image corresponds to the average of 150 pictures. **(b)** AFM images of 5-day-old fibrils of the three H46 variants. White squares indicate the zoom region displayed in panels below.
